# Representational similarity learning reveals a graded multidimensional semantic space in the human anterior temporal cortex

**DOI:** 10.1101/2022.10.27.514039

**Authors:** Christopher R. Cox, Timothy T. Rogers, Akihiro Shimotake, Takayuki Kikuchi, Takeharu Kunieda, Susumu Miyamoto, Ryosuke Takahashi, Riki Matsumoto, Akio Ikeda, Matthew A. Lambon Ralph

## Abstract

Neurocognitive models of semantic memory have proposed that the ventral anterior temporal lobes (vATLs) encode a graded and multidimensional semantic space—yet neuroimaging studies seeking brain regions that encode semantic structure rarely identify these areas. In simulations we show that this discrepancy may arise from a crucial mismatch between theory and analysis approach. Utilizing an analysis recently formulated to investigate graded multidimensional representations, *representational similarity learning* (RSL), we decoded semantic structure from ECoG data collected from the vATL cortical surface while participants named line drawings of common items. The results reveal a graded, multidimensional semantic space encoded in neural activity across the vATL, which evolves over time and simultaneously expresses both broad and finer-grained semantic structure amongst animate and inanimate concepts. The work resolves the apparent discrepancy within the semantic cognition literature and, more importantly, suggests a new approach to discovering representational structure in neural data more generally.

**Conflict of interest:** The Department of Epilepsy, Movement Disorders, and Physiology, Kyoto University Graduate School of Medicine conducts Industry-Academia Collaboration Courses, supported by Eisai Co., Ltd, Nihon Kohden Corporation, Otsuka Pharmaceutical Co., and UCB Japan Co., Ltd.

## Introduction

If you encounter a wolf when walking home through a dark wood, your mind readily accomplishes some remarkable feats: it anticipates the animal’s likely behavior, perhaps slinking closer toward you; it assigns the thing a name, which you might shout to alert others (“wolf!”); and it directs you to change your own plans, maybe running back down the path. These feats arise from the human ability to discern conceptual structure—to realize that the wolf, despite its resemblance to friendly dogs in town, is nevertheless a quite different sort of animal.

This ability to represent and exploit conceptual structure is central to human semantic cognition. Such structure is *graded* in that similarities vary along a continuum: wolves are highly similar to coyotes, partially similar to elk, and quite distinct from birch trees or canoes. It is also *multidimensional* in that concepts vary along a myriad of independent components: wolves and dogs are similar in their shapes, parts, movements, and phylogeny, but different in their behaviors, habitats, and diets. To capture these properties of knowledge, computational approaches to semantics often represent concepts with *vector spaces*: the meaning of a word or image is expressed as a point in an *n*-dimensional space (or equivalently as an *n-*dimensional vector) such that the proximity between points expresses the similarity in meaning between the denoted concepts. The dimensions of the vector need not correspond to nameable conceptual components like *habitat* or *diet*; instead they may define a space, with different concepts corresponding to different points in the space, and with the distances between points capturing the degree of semantic/conceptual relatedness between concepts (Frisby et al., 2023). Cognitive science and machine-learning offer many techniques for estimating semantic vector spaces from natural language (Pereira et al., 2016), feature norms (McRae et al., 2005), or similarity-judgments (Hebart et al., 2020), and these methods have provided a critical empirical foundation for studying human conceptual knowledge.

The well-known “hub and spokes” theory of semantic representation suggests that the anterior temporal lobes (ATLs) encode a graded, multidimensional semantic vector space that expresses conceptual similarity structure for all concepts, extracted across all input and output modalities, and from our experiences of each concept across time (Jackson et al., 2021; Lambon Ralph et al., 2017; Patterson et al., 2007). This proposal has been useful for understanding patterns of semantic deficits arising from temporal lobe pathology in fronto-temporal dementia (Acosta-Cabronero et al., 2011; Hodges & Patterson, 2007; Lambon Ralph, 2014; Lambon Ralph et al., 2010; Rogers et al., 2004; Snowden et al., 1989), anterior temporal resection (Drane et al., 2008; Lambon Ralph et al., 2010; Lambon Ralph et al., 2012; Rice et al., 2018; Schapiro et al., 2013), and herpes viral encephalitis (Gainotti, 2018; Lambon Ralph et al., 2007; Noppeney et al., 2007); stimulation and evoked response direct neurophysiological explorations (Abel et al., 2015; Y. Chen et al., 2016; Shimotake et al., 2015); the effects of transcranial magnetic stimulation in ATL and other parts of the cortical semantic system (Binney et al., 2010; Binney & Lambon Ralph, 2015; Jefferies, 2013; Lambon Ralph et al., 2009; Pobric et al., 2007, 2010); and a variety of behavioral phenomena in developing and mature cognition (Chen et al., 2017; Jackson et al., 2021; Rogers & McClelland, 2004, 2014).

Yet direct empirical tests of this proposal—representational similarity analysis (RSA) of functional imaging data collected while people perform semantic tasks on words or pictures—have not generally tended to support it. A recent review identifies 24 studies applying RSA to uncover semantic representations in the brain (Frisby et al., 2023); of these, 18 (75%) failed to identify semantic structure in the anterior temporal cortex (for the exceptions, see Bruffaerts et al., 2013; Devereux et al., 2018; Fairhall & Caramazza, 2013; Martin et al., 2018; Peelen & Caramazza, 2012). Many of these studies instead find that semantic structure is encoded in brain areas not otherwise thought to be critical to semantic representation, including posterior cortical regions (Connolly et al., 2012), inferior and superior frontal and motor cortex (Carota et al., 2017; Wang et al., 2017), the left pars triangularis (Liuzzi et al., 2017), right superior parietal cortex (Wang et al., 2017), the insula and occipeto-parietal cortex (Kivisaari et al., 2019), and the posterior cingulate cortex (Fairhall & Caramazza, 2013). Thus RSA studies often yield results that seem puzzling given the broader literature, finding that semantic structure is encoded in many areas throughout cortex but not in the ATL.

The frequent failure of RSA to find semantic structure in ATL may reflect limitations of fMRI, which, without specialized acquisition protocols, can yield poor signal in ventral aspects of this brain area (Halai et al., 2015; Halai et al., 2014). An important study by (Y. Chen et al., 2016) suggests, however, that this is not the full story. The authors collected intracranial grid electrode voltages (ECoG) from the surface of ventral anterior temporal lobes (vATL) while participants named line drawings of familiar items, then conducted a searchlight-based semantic RSA from these data. Consistent with the semantic-hub model, they found an anterior fusiform area where similarities in the evoked neural response correlated significantly with the target semantic similarities. Critically, however, they further showed that the evoked neural similarities correlated *equally well* with a binary target matrix that only encoded whether a stimulus was animate or inanimate. This result is consistent with an alternative view that, while ECoG in vATL may express a coarse binary animacy distinction, it does *not* otherwise encode graded or multidimensional semantic structure within or between these domains. ECoG is not affected by the magnetic field inhomogeneities that affect fMRI in vATL, so the finding is not easily attributable to poor signal or other data artifacts.

Motivated by these observations, this paper considers an alternative hypothesis about the discrepant findings for RSA versus the broader literature: that it arises because RSA, as typically practiced, is not well-suited to finding *<u>graded multidimensional</u>* vector spaces of the kind the ATL is hypothesized to encode. This may seem surprising, since RSA was developed specifically as a tool for finding cognitive similarity structure in neuroimaging data (Kriegeskorte, Mur, & Bandettini, 2008)—yet, as we will demonstrate, the reliance of the approach on correlation significantly limits the kinds of signal it can detect.

Study 1 considers how RSA constrains what can be discovered about neuro-semantic representations, by applying the approach to simulated data where the target signals are designed to encode elements of structure in a true semantic similarity matrix derived from feature-listing data. The results illustrate how and why RSA can both miss real signal and produce positive results with the potential to mislead. Study 2 then introduces a different approach to neural decoding, *representational similarity learning* (RSL; Oswal et al., 2016), that can remediate these issues by making explicit which aspects of the target structure can be successfully decoded, and by operationalizing hypotheses about the neurosemantic code directly within the decoding model. Study 3 extends the RSL approach to the analysis of very large neural datasets and applies it to investigate graded multidimensional semantic structure in ECoG data recorded from the surface of human vATL—the same data for which RSA data failed to detect graded, multidimensional and cross-domain semantic structure in a prior study (Y. Chen et al., 2016). We empirically compare results yielded by RSL vs. RSA on these data, with results that resolve the contradiction in the literature and suggest a new path for multivariate neural decoding more generally.

### Simulation study 1: Evaluating RSA

Simulation 1 evaluated whether the results yielded by RSA reliably indicate whether a set of neural measurements encode graded, multidimensional semantic similarity structure of the kind typically sought in functional imaging studies. By *graded*, we mean that the measurements encode varying and continuous degrees of similarity between items, rather than discrete or categorical distinctions between items. By *multidimensional*, we mean that the measurements encode variation along more than one orthogonal component of a representational space. Though the general approach is well known, its limitations may be less familiar, so we begin with a brief overview of the method and some of the challenges it faces when seeking graded, multivariate structure.

RSA aims to find sets of neural features—voxels, electrodes, or other measurements of neuro-physiological activity—whose responses to various cognitive events (e.g., the perception of a stimulus) jointly encode an independently-measured target structure. Typically, the target structure is a *representational similarity matrix* (RSM; sometimes called a representational dissimilarity matrix or RDM) in which the rows and columns correspond to the different stimuli in an experiment and the entries indicate the cognitive/representational similarities between stimuli. For semantic representation, entries in the target RSM indicate similarity of meaning amongst pairs of concepts as estimated from behavioral or corpus data. To determine if responses in a set of neural features (e.g., voxels, electrodes, EEG sources, etc.) encode the target similarities, the experimenter estimates the pattern of neural activity evoked over features by each stimulus and computes the pairwise similarities between these to create a *neural similarity matrix* (NSM). Correlations between the RSM and NSM are computed separately for each participant, and brain regions where these are reliably greater than zero are interpreted as encoding the target structure.

This approach faces at least five challenges when used to find *graded*, *multidimensional* structure in neural codes:

1. *Discrete vs graded structure*: reliable correlations with a continuous-valued target RSM can arise even if the neural response is discrete or categorical. Thus, a positive result does not indicate that the neural response encodes *graded* structure even if such structure is present in the target RSM.
2. *Unidimensionality*: the correlation between RSM and NSM is inherently unidimensional. Thus, a positive result on its own does not provide evidence of multidimensional structure in the neural response.
3. *A priori feature selection*: the experimenter must decide ahead of time which neural measurements to use when computing the NSM (e.g. an ROI or a searchlight). If the informative features fall across different ROIs or searchlights or are intermixed with many non-informative features within an ROI/searchlight, the approach can fail to discover them.
4. *Equal importance*: all selected neural features are equally weighted when computing the NSM. If the informative neural features are sparse or vary in signal strength, equal weighting of all features may lead to a null result. Thus, a null result does not indicate that the selected neural features carry no target information.
5. *Masked dimensionality*: reliable correlations between RSM and NSM can be driven by a subset of the RSM structure or items, or even by one or two extreme values. Thus, a positive result does not imply that all components of the target RSM are encoded.

Simulation 1 assessed how these characteristics of RSA will influence discovery of neuro-semantic representations by applying the approach to synthetic data designed to encode different aspects of a real semantic similarity matrix.

## Methods

The target RSM was computed from semantic feature norms for 100 items, half animate and half inanimate, collected in prior work (Dilkina & Lambon Ralph, 2013). The feature vectors for each item were aggregated in a matrix with rows indicating words, columns indicating features, and binary entries indicating whether the referent of the word possesses that feature. Matrix columns were mean-centered and the RSM was computed as the cosine similarity for all pairs of row vectors.

To understand the latent structure of the RSM, we used singular value decomposition (SVD) to extract three components accounting for 90% of the variance in the full matrix (81.1%, 4.4%, 4.0%, respectively). SVD is a common matrix decomposition algorithm that reduces a matrix to a small number of orthogonal components ordered by the proportion of variance they explain. After applying it to the RSM, each of the 100 items in the set receives a coordinate position along the three orthogonal components.

Figure 1 shows the coordinates for all 100 items on each latent dimension, color coded by semantic category (see Appendix A for full list of items and how they were categorized). The decomposition reveals graded semantic structure along each dimension. The first separates animate from inanimate items but also individuates subcategories in each domain. The second strongly differentiates subcategories of animals but also weakly distinguishes inanimate subcategories. The third differentiates the inanimate items, though the resulting spread is less “clumpy” than within animate items as commonly found with such data (Lambon Ralph et al., 2007). From the SVD it is clear that the semantic vector space is both multidimensional and graded by the definitions offered earlier.

**Figure 1.**
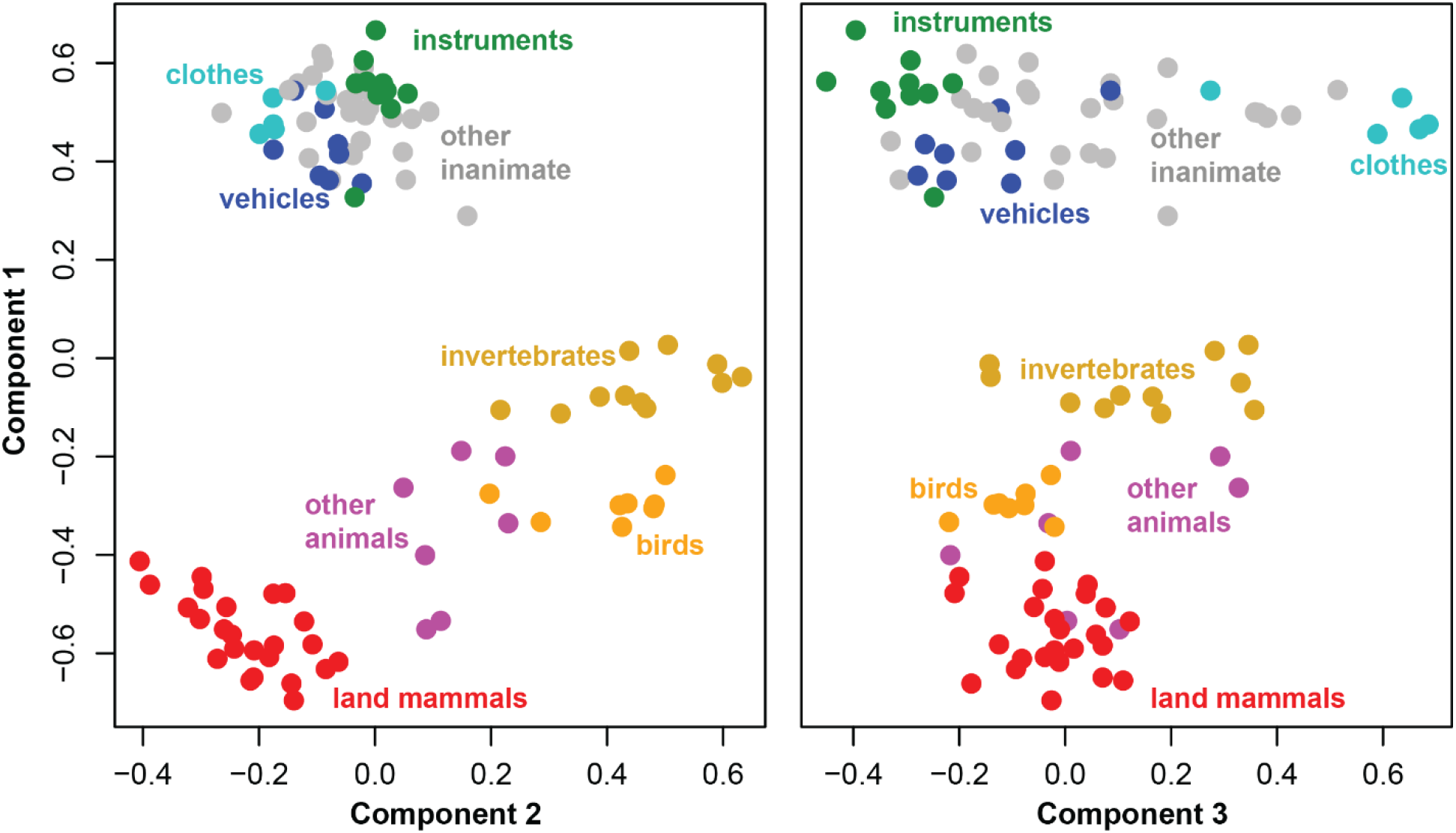
Coordinates of the 100 stimuli in the three-dimensional singular-value-decomposition of the semantic similarity matrix. Colors show category membership within each domain. Component 1 separates animate from inanimate items, but also separates animate sub-categories. Component 2 largely separates animate sub-categories, while component 3 largely separates inanimate sub-categories.

The simulation assessed what results RSA would yield when applied to a set of signals designed to encode elements of this underlying semantic structure, when those signals are perturbed by noise and embedded in a population together with other signals that do not code semantic information (as in brain imaging data). Specifically, we created five simulated datasets, each capturing a different aspect of semantic structure in the target RSM. Each dataset contained simulated responses of 24 features to the 100 items, as a simple model analog of the responses of, for instance, voxels within an ROI. To capture the fact that a given ROI may contain both signal-carrying and uninformative voxels, half of the features encoded true semantic information about the target items while half adopted random values sampled from a uniform distribution in the same range.

The five datasets varied in which aspects of the target matrix they encoded, as illustrated in the schematic plots in Figure 2A. In the *binary condition*, all signal-carrying features adopted one state (-1 or 1) for animate items and the opposite state for inanimate items—thus the code was both unidimensional and discrete (i.e., non-graded). In three *one-D conditions*, all signal-carrying features only encoded a single latent component of the target RSM—either the first, second, or third singular vector as shown in Figure 1. Thus, each encoded *graded* similarity structure (the distances between items were continuous, not binary/discrete) but only *one* dimension of variation. In the *full structure condition,* the signal-carrying features jointly encoded all three latent components of variation in the target RMS, with four features dedicated to each of the three components. Thus, the code was both graded and multidimensional.

**Figure 2.**
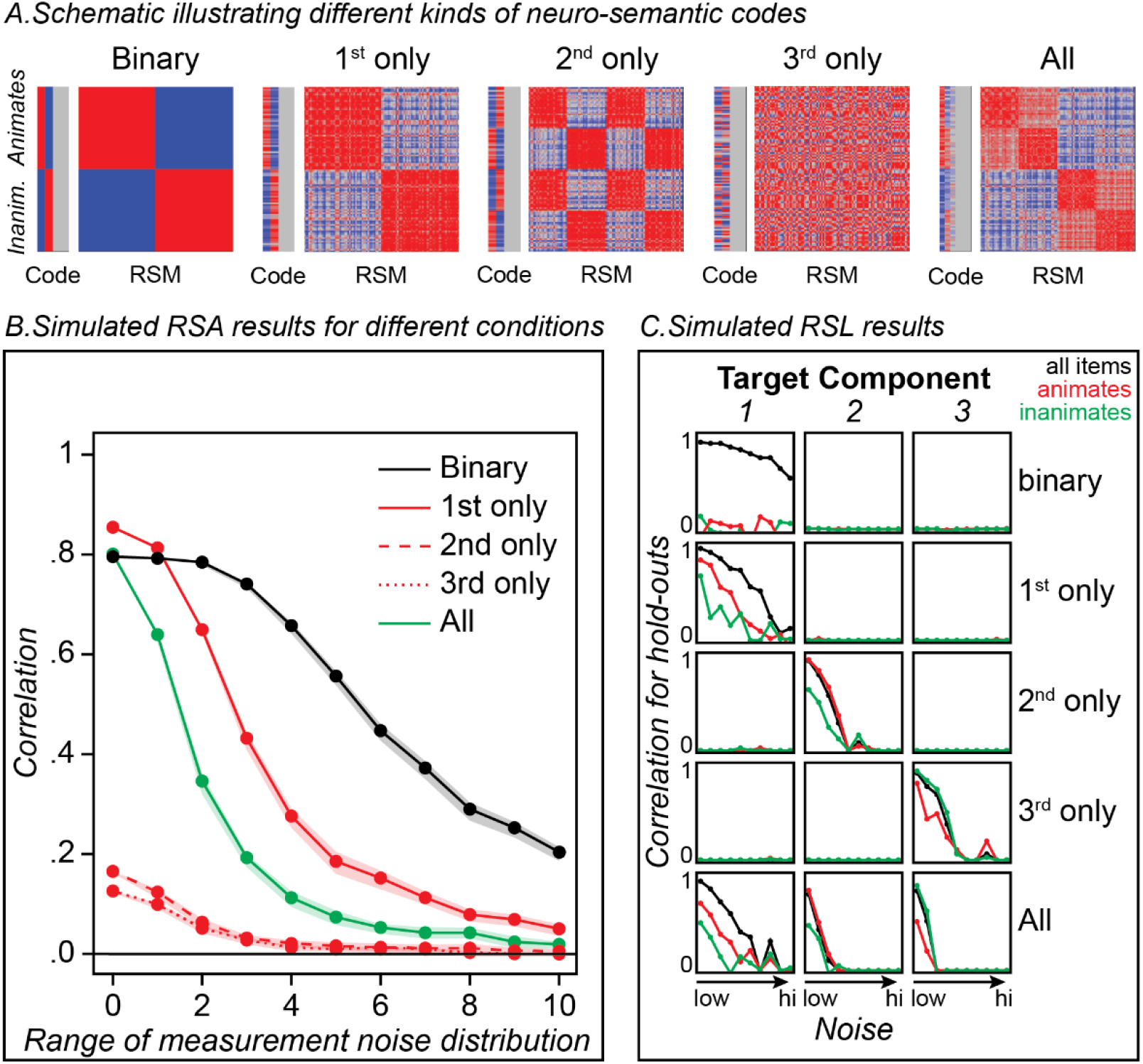
Simulation results. A. Each plot illustrates different ways that a neural response might encode some aspects of a target RSM, schematized in the rightmost plot (“All”). Vertical bars show the hypothesized responses of neural features to various animate and inanimate stimuli (red more active; blue less active; gray random) while the squares show the neural similarity matrices that would then result (red high similarity, blue low). All conditions encode some components of the full RSM and so should be detected by a multivariate method seeking semantic structure. B. Curves showing the expected results of standard RSA when used to decode real semantic structure from the target RSM under different hypotheses about the neural code and increasing amounts of measurement noise. Dots show the mean correlation while ribbons show the 95% confidence interval. The approach can yield robust results when the neural signal is discrete and/or unidimensional, even under substantial noise, and can yield weak or null results when the neural signal faithfully encodes weaker components of the RSM, even under low-noise conditions. C. Results of RSL applied to the same simulated data, showing the correlation between true and predicted coordinates for held-out items on each latent component of the target matrix (columns) at the same 11 levels of noise as in the RSA, for the five different simulation conditions (rows), and computed across all items (black lines), animate items (red), or inanimate items (green). RSL shows a positive result in each case, identifies which components of the matrix are present in the neural code, and reveals reliable within-domain decoding only when the neural code expresses continuous similarity structure.

For each signal condition, we distorted the response of each feature to each stimulus with measurement noise sampled independently from a uniform distribution centered on 0, generated a simulated NSM, then computed the correlation between the simulated NSM and target semantic RSM. The varying levels of noise allowed us to investigate RSA behavior across a wide range of signal-to-noise ratios. This procedure was conducted 20 times at each of 11 increasing noise levels. We then computed, across runs at each noise level, the mean and 95% confidence intervals of the Pearson correlation between simulated NSM and RSM.

## Results

The results are shown in Figure 2B. The binary code showed robust correlations across the full noise-spectrum, illustrating that RSA with a graded and multidimensional target RSM can yield a positive result even when the neural code is neither continuous nor multidimensional. Amongst the one-D codes, RSA showed a strong positive result if the simulated neural response encoded the first singular vector of the target matrix, but much weaker results that decayed to zero with increasing noise when the neural response encoded other components. The weak results for components 2 and 3 arise simply because the raw similarities encoded in the target RSM are most strongly determined by the first component. Simulated NSMs that encode just the second or third component thus express similarities that do not correlate strongly with the pairwise similarities encoded in the target matrix. Thus, RSA can yield weak/null results even when the features under consideration *do* reliably encode semantic information that is orthogonal to the primary component of the RSM. Throughout the noise range assessed, RSA yielded the most reliable results when the simulated features encoded a binary domain distinction or just the first component of variation. For these scenarios, all signal-carrying features encode the same information, so noise “cancels out” across features and the signal remains robust. When the same number of features encode three orthogonal components, fewer resources are dedicated to each, and corruption from noise more seriously degrades the signal—so that, counter-intuitively, the approach yielded less reliable results when the simulated features encode all three dimensions of the target matrix.

## Discussion

The simulation shows that, when decoding semantic similarity structure of the kind captured by semantic feature norms, RSA can yield a positive result when the neural response is discrete and/or unidimensional, *even* though the target matrix is continuous and multidimensional. Additionally, it can yield weak or null results when the neural response *does* encode underlying dimensions of the target matrix beyond the strongest. Thus, the simulation serves as a demonstration that RSA, as typically deployed, may not reliably indicate whether a set of neural signals encode the graded, multidimensional semantic structure existing in a target RSM. Of course, the specific pattern of results obtained will depend upon the ratio of informative to uninformative features within the selected set and the amount of noise perturbing the signal, in addition to the structure encoded by the informative features.

One response to these challenges may be to consider the degree of correlation between a given NSM and multiple different RSMs, each encoding a different kind of structure. Yet simulation 1 suggests that this approach remains limited. Recall that Y. Chen et al. (2016) found that an ECoG-based NSM correlated equally well with an animacy-based binary similarity matrix as with a full continuous and multidimensional matrix. The rationale for the comparison was the expectation that, if the NSM encodes the full similarity structure, it should show a higher correlation with the target semantic matrix than with the binary matrix. Figure 1B shows that this is not necessarily so—in fact the binary code showed as good or better correlation with the target semantic matrix than did the full (continuous, multidimensional) code. Thus, the comparison of fits with the two different target matrices did not help to resolve the nature of the underlying neural code.

Or, consider the interesting findings of Clarke and Tyler (2014), who used RSA to seek areas encoding semantic structure both across and within superordinate domains (such as “animals” and “vehicles”). When the target matrix included all items from all domains, the analysis identified the vATL as encoding semantic structure. When limited only to items within one semantic domain, however, the vATL did not appear to encode such structure. The finding could mean that vATL only encodes discrete distinctions between superordinate semantic domains, analogous to the binary condition in the simulation—but the same result could also obtain even if the vATL does encode within-domain structure, since such structure is mainly encoded, within the semantic target matrix, by components orthogonal to the primary dimension.

More sophisticated use of RSA may overcome some of these difficulties in interpretation; for instance, the researcher might carefully select stimulus items to ensure that multiple orthogonal components of a representational space are equally strongly expressed in the RSM, or to ensure that different target representational spaces are maximally differentiated by the selected stimulus items. Indeed, one of the few RSA studies to report semantic representational structure in ATL adopted this approach, ensuring that semantic and visual similarity structure were completely deconfounded in their stimuli and then comparing results for these two differently-structure target matrices (Martin et al., 2018). While such studies are elegant and informative, they are also challenging to design and may prevent researchers from exploring fuller or more naturalistic distributions of stimuli. For these reasons, we were motivated to consider an alternative approach to discovery of graded multidimensional representations in neural data.

### Simulation study 2: Decoding with RSL

The approach we developed is called *representational similarity learning* (RSL; Oswal et al., 2016). Rather than using pairwise similarities in the RSM as the target values for correlation, RSL first decomposes the matrix into orthogonal latent components, effectively re-representing each stimulus item as a point in a low-dimensional semantic space as shown in Figure 1. It then uses linear regression to fit *decoding models* that predict the coordinates of each item along each dimension from their evoked neural responses. Simulation 2 assessed whether this approach can reveal the graded and/or multidimensional structure obscured by RSA.

## Methods

Using the same simulated neural signals from Simulation 1 as predictors, we fit ordinary least-squares (OLS) regression models to predict coordinates of each stimulus along each of the three latent semantic dimensions shown in Figure 1. For each simulated feature set and each of 11 increasing levels of uniformly distributed noise (i.e., the same noise manipulation as for Simulation 1), each example was assigned to one of ten mutually exclusive test sets such that each test set had 5 animate and 5 inanimate items. Holding out each test set in turn, regression models were fit to the remaining 90% of the examples (the training set). The model that was fit to the training set predicts coordinates for the 10% of examples withheld as the test set. After iterating over the ten test sets, a hold-out prediction exists for every item. These were concatenated and correlated with the target embedding one dimension at a time. This process was repeated 20 times with different samples of noise. Hold-out accuracy for each condition and noise level was estimated as the mean over simulation repetitions.

Multidimensional structure in the neural code should be revealed by reliable prediction of coordinates along more the one latent dimension. To assess whether the neural code captures graded similarity, we evaluated how well the fitted model predicted coordinates amongst just the animate items or just the inanimate items. If the neural signal only categorically differentiates animate from inanimate stimuli, then predicted and true values should correlate significantly when models are evaluated on all items but should *not* correlate amongst just the animate or just the inanimate items considered separately. Reliable intra- and inter-domain correlations between true and predicted values thus indicate that the neural code captures a graded degree of similarity structure.

## Results

Figure 2C shows the mean correlation between true and predicted coordinates for held-out items on each of the three target components (columns), for the five different neural coding scenarios (rows), and computed across all items (black lines), animate items (red), or inanimate items (green). The pattern of results reveals the information encoded in each simulated neural response. With the binary neural signal (top row), predicted and true coordinates correlated strongly for models fit to all items but not separately for animates vs. inanimates. With continuous one-D signals, strong correlations were observed for models fit to all items and within each domain separately, but only for the dimension encoded by the neural response. When the neural signal encoded graded multidimensional structure, reliable correlations were observed on all three latent components, for models fit to all items and fit separately to each domain. Thus, in simulation, the pattern of results uniquely revealed whether the neural signal encodes graded or discrete structure, and which semantic dimensions it captures.

## Discussion

In contrast to the RSA simulation, RSL yielded a different, diagnostic pattern of results for each condition, indicating which aspects of the semantic target matrix were encoded within the simulated neural responses. The discreteness of the binary code was revealed by reliable prediction across domains and no reliable prediction within domain. In the 1D neural-structure conditions, RSL showed reliable prediction only on the latent dimensions encoded by the simulated neural signal, and not on the other dimensions. Reliable decoding both within and between domains in such conditions indicated, in contrast to the binary condition, that the signal is graded rather than discrete. When the simulated neural signal encoded all three semantic dimensions, RSL showed reliable decoding of each. Thus, by considering decoding accuracy on orthogonal components of the target matrix, RSL can test whether the neural code expresses multidimensional structure; and by considering decoding accuracy separately for different semantic domains, it can test whether the code is graded.

### Study 3: Decoding semantic structure from ECoG using RSL

Study 3 evaluated whether neural signals in vATL encode a graded, multidimensional and domain-general semantic space by using RSL to decode semantic structure in ECoG data collected in a prior study of object naming (which employed the same 100 items used in the simulations; Y. Chen et al., 2016). This dataset is especially useful in the current context for two reasons.

First, ECoG avoids some of the limitations of non-invasive imaging methods that can make structure in vATL difficult to decode. One recent study suggested that the neuro-semantic code for animacy in vATL changes rapidly and nonlinearly over the course of stimulus processing so that techniques sacrificing either spatial or temporal resolution, including EEG, MEG, and fMRI, may obscure important signal (Rogers et al., 2021). Additionally, magnetic field inhomogeneities make it difficult to resolve clear BOLD signal in vATL without special acquisition sequences (Halai et al., 2015; Halai et al., 2014). ECoG provides much better spatial and temporal resolution of neurophysiological signals at the surface of cortex and does not suffer from the signal-detection issues that challenge fMRI, so any failure to discover fine-grained / multivariate semantic structure in the data cannot reflect these limitations.

Second, the results of the original RSA study (Y. Chen et al., 2016) suggest that the measured signals, though reliably differentiating animate from inanimate items, do *not* encode finer grained or multidimensional semantic structure. Specifically, the authors compared the correlation of the stimulus evoked NSM to two different target matrices: one encoding the full graded and multidimensional semantic similarities amongst stimuli, and a second encoding just the binary distinction between animate vs inanimate items. If the vATLs encode graded multidimensional semantic structure, one might expect the NSM to correlate more strongly with the full target matrix than with the binary animacy matrix. Instead, the authors found statistically significant and *equally strong* correlations with both target matrices. One interpretation of this finding is that the measured neural signals do *not* in fact encode graded or multidimensional semantic structure, but only serve to discretely differentiate animate and inanimate items—a conclusion also consistent with Clarke & Tyler’s (2014) RSA results described earlier. The data thus allow us to test an alternative hypothesis: that the measured signals *do* encode a graded, multidimensional and domain-general semantic space, in a manner that is invisible to RSA. If so, RSL should reliably decode variation along multiple orthogonal components of the target semantic matrix, considering all items together as well as within animate and inanimate domains considered separately.

Applying RSL to ECoG or any other brain imaging data, however, faces an immediate challenge: such technologies produce many more spatiotemporal neural measurements than there are stimuli, so regression models predicting stimulus characteristics from neural data are ill-defined (i.e., there exist infinite solutions that can perfectly predict outcomes on training data even from random input data). To find a unique fit without a priori feature selection, the regression must be *regularized* to satisfy some additional constraint that will guarantee a single unique solution for model fitting. For instance, the weight optimization might jointly minimize prediction error and the size of the model coefficients, measured as the sum of their absolute values (the *L1-norm*). Since this sum can be minimized by placing zero coefficients on many predictors, this form of regularization promotes a sparse solution in which only a few predictors have non-zero coefficients in the final model (e.g., Rogers et al., 2021). Other regularizers (such as the L2 norm or the elastic net) enforce different constraints on model fitting, and thus lead to other solutions when applied to the same dataset (Cox & Rogers, 2021).

In this sense, the selection of a regularization function amounts to a prior hypothesis stipulating how the neural signal is likely to be structured. As recently argued in (Frisby et al., 2023), the choice of regularizer should be informed by explicit hypotheses about the nature of the signal to be decoded—in this case, a hypothesis about how neurophysiological signals measured by surface electrodes in the brain might encode a graded, multi-variate semantic space.

To that end, we adopted a regularization function called the *group-ordered-weighted LASSO* (grOWL) loss (Oswal et al., 2016) that was designed in prior work to capture three theoretically-motivated assumptions about the structure of neural representations. First, it assumes the signal is *sparse*: of all measurements taken, only a relatively small proportion encode target information of interest, so the regularizer should place zero coefficients on many predictors (as with L1 regularization). Second, it assumes *redundancy* in the true signal: neural populations that do encode target information will be correlated in their activity patterns over items, so the regularizer should spread similar coefficients across correlated signal-carrying populations (similar to L2 regularization). Third, it assumes that signal-carrying populations are unlikely to be “axis-aligned” with the target semantic space (i.e., responding only to variation along a single semantic dimension) but more likely encode multiple, varying “directions” within the space. That is, each neural population likely carries information about an item’s coordinates along multiple dimensions of the target space. We will refer to this as the *spanning* assumption, since the selected neural features span the target representational space but are unlikely to only encode distinct, individual dimensions. To enforce this assumption, the regularizer should prefer solutions where coefficients on a given neural feature are either all zero (the feature does not carry any information) or all non-zero (it predicts some variation along all target dimensions).

Oswal et al. (2016) showed that all three assumptions can be captured in a single convex loss function, and also provided analytic guarantees and an example application on an fMRI dataset. We describe the logic briefly here; Appendix B provides a formal description of the decoding model and grOWL regularizer, including a definition of the loss and explanation of how it encourages these properties in the decoding matrix.

Rather than fitting a separate regression model for each latent component of the target RSM (as in the simulation), we instead predict all latent components (Figure 3A) simultaneously by estimating the parameters of a *decoding matrix* β (Figure 3C) with regularized multitask regression. Suppose the coordinates of *n* stimuli on *r* latent semantic dimensions are stored in matrix ***U***_*n*×*r*_ while the responses of *m* neural features to the stimuli are stored in matrix ***X***_*n*×*m*_. RSL models the entries in ***U*** as the matrix product of the neural activations in ***X*** and the decoding matrix **β**_*m*×*r*_: ***U*** = ***X*β**. Each row of **β** corresponds to one neural feature (voxel, electrode, etc.), and each column encodes weights for each neural feature when predicting stimulus coordinates on the corresponding dimension of the target matrix ***U***. Thus, the assumptions about neural signal just listed can be formalized as constraints on the structure of the decoding matrix **β** (Figure 3D). To capture the sparsity and spanning assumptions, **β** is constrained to be *row-sparse*: most rows have all zero values (sparsity), and the rest have all non-zero values (spanning). To capture the redundancy assumption, highly correlated and signal-carrying neural features receive similar row-vectors in **β** (Figure 3D). The precise degree of sparsity and redundancy in the decoding matrix are controlled by hyper-parameters that are tuned via cross-validation. Figure 3 shows the full workflow decoding semantic structure from ECoG data.

**Figure 3.**
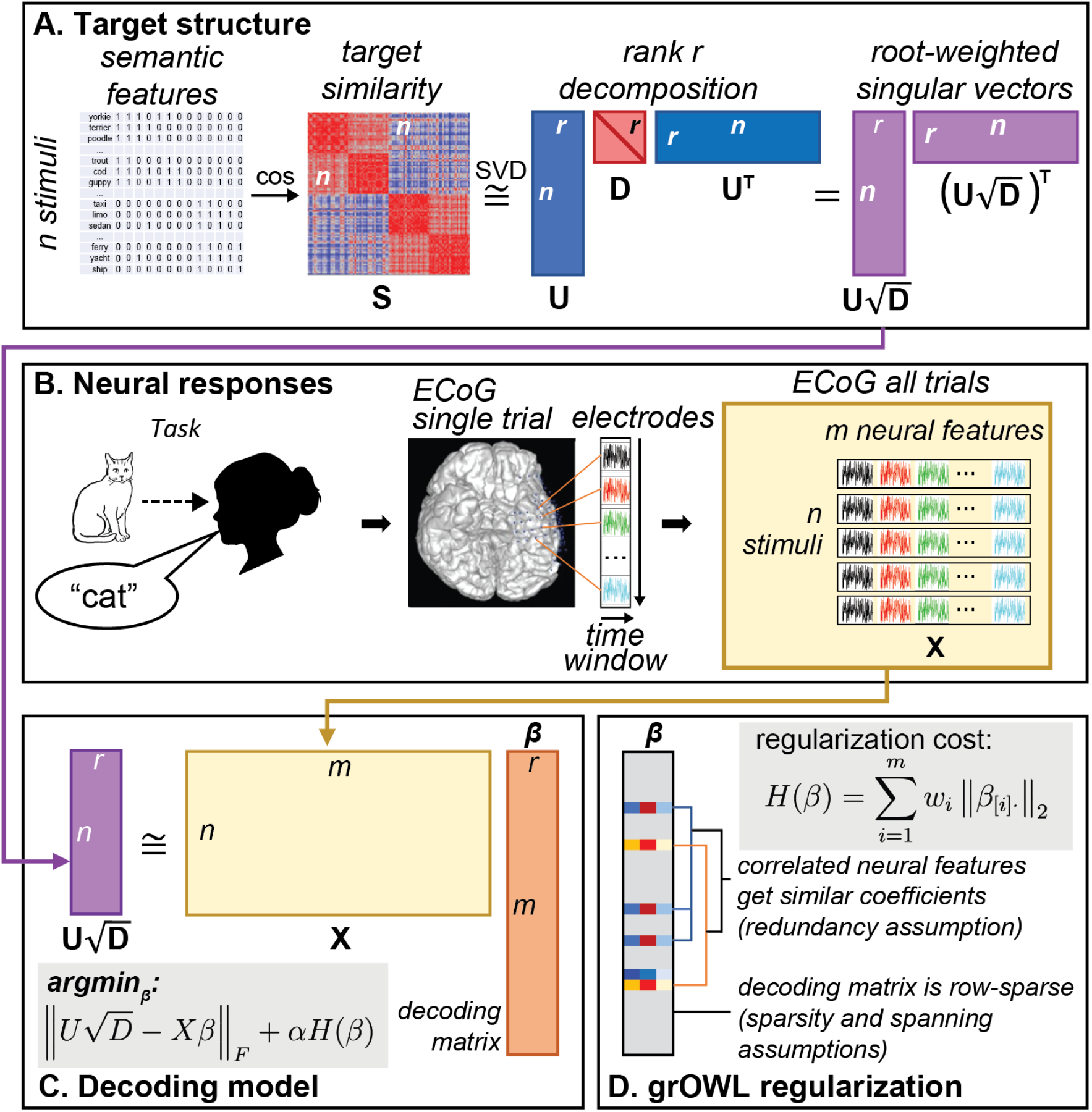
Representational similarity learning. A. To generate the target structure for decoding, semantic feature vectors for each of *n* items are mean-centered and converted to a cosine similarity matrix ***S***_*n*×*n*_ of approximate rank *r*. This is decomposed into matrix ***U***_*n*×*r*_ containing *r* orthogonal singular vectors and diagonal matrix ***D***_*r*×*r*_ containing singular values for each vector. The product 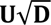 is an *nxr* matrix whose columns contain *r* root-weighted singular vectors that are the target of the decoding model. Note that the product of this matrix with its transpose provides an estimate of the original matrix ***S***_*n*×*n*_. B. Neural responses in a picture-naming task are recorded for each stimulus as intra-cranial voltage potentials collected at 1000Hz over electrodes implanted in vATL. Data are concatenated across electrodes to create a single *m*-dimensional “neural feature vector” for each stimulus, where *m* is the total number of electrodes times the size of the decoding window in milliseconds. All such vectors are compiled into a neural feature matrix ***X***_*n*×*m*_ which contains the predictors for the decoding model. C. The target structure 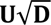 is then modeled as the product of the neural feature matrix ***X*** and a decoding matrix of regression coefficients **β** whose values are chosen via gradient descent to minimize prediction error plus the regularization penalty *H*(**β**) scaled by the weighting parameter α. D. Definition of the group-ordered-weighted LASSO (grOWL) regularization function and illustration of the structure it promotes in the decoding matrix.

With this overview we can consider how RSL can be applied to assess whether neural data encode a multidimensional and graded similarity space. Parameters in **β** are estimated for a batch of training data using regression regularized with the grOWL loss. The decoding model is then applied to predict the coordinates of held-out items along each dimension in the target representation space. If predicted and true coordinates for held-out items correlate reliably on more than one dimension, the neural signal must encode multidimensional structure. To assess whether the neural code captures graded similarity, we additionally evaluate the regression models separately for animate and inanimate items, as in the simulation. If the neural signal only categorically differentiates animate from inanimate stimuli as suggested by the prior RSA analysis in Y. Chen et al. (2016), then predicted and true values should correlate significantly across all items, but not for animate or inanimate subsets considered separately. Observation of both inter- and intra-domain correlations thus indicates that the neural code captures a graded degree of similarity structure. Finally, to assess how well the decoder recovers the target semantic similarities, the predicted coordinates for held-out items on all three latent dimensions can be used to estimate semantic distances between stimulus pairs. These estimates can then be correlated with true semantic distances, similar to standard RSA.

## Methods

We applied this approach to the same ECoG dataset where prior work using RSA found no evidence for graded multidimensional structure (Y. Chen et al., 2016). The study was approved by the ethics committee of the Kyoto University Graduate School of Medicine (No. C533) and participants provided written informed consent. The dataset contains voltages measured from platinum subdural grid electrodes (inter-electrode distance 1 cm and recording diameter 2.3mm; Ad-Tech, WI) implanted in the surface of left (8) or right (2) vATL in ten patients undergoing preparation for intractable epilepsy (9) or brain tumor (1) while they named line drawings depicting the 100 items described in Study 1. Electrodes were implanted in the right hemisphere for two patients where WADA testing did not clearly indicate left-lateralization of language.

Each patient had between 6 and 32 electrodes (mean of 20) covering vATL. All patients with epilepsy had seizure onset zones outside the anterior fusiform region, except one patient for whom it was not possible to localize the core seizure onset region. Data were sampled at 1000Hz (eight patients) or 2000Hz (two patients) with a band-pass filter of 0.016 – 300 (eight patients) or 0.016 – 600Hz (two patients). The dataset included, for each stimulus, voltages measured at each electrode over a 1s window from stimulus onset and downsampled to 100Hz. Thus, for a participant with 20 electrodes, each stimulus was associated with a 2,000-element vector of voltages (20 electrodes x 100 time points). The ECoG data were preprocessed exactly as in Y. Chen et al (2016).

### Stimuli and Procedure

One hundred line-drawings (50 animate and 50 inanimate items) were obtained from previous norming studies (Morrison et al., 1997; Snodgrass & Vanderwart, 1980). See Y. Chen et al. (2016) for a complete list. Animate and inanimate stimuli were matched on age of acquisition, visual complexity, familiarity, and word frequency, and had high name agreement. Each stimulus was presented on a computer screen for 5 seconds, one after another with no interstimulus interval, once during each of four sessions (four times in total). Each session proceeded in a different random order. Participants were instructed to name each item as quickly and accurately as possible. Participants were video and audio recorded during the experiment. Video was used to monitor eye fixations and general attention to the task. The mean response latency was 1190ms.

### Semantic similarity structure and dimensionality reduction

The same target RSM and 3-dimensional embedding used in the simulated analysis are reused for the ECoG analysis.

#### ECoG data preprocessing

Preprocessing was performed in MATLAB. Data were downsampled to 100Hz by averaging measurements within 10ms boxcars. A prior study applying pattern classification to these data found near identical results when analyzing raw voltages versus voltages referenced to the electrode beneath the galea aponeurotica or to the scalp electrode on the mastoid process contralateral to the side of electrode implantation (Rogers et al., 2021). For this reason, and because the location of the reference electrode varied across patients, we analyzed all voltages without referencing. For each stimulus we retained data for 1000ms from stimulus onset. While the trial epoch may include the onset of articulation toward the end, the critical results cannot reflect such motor activity since all key phenomena are observed early in the epoch. Baseline correction was not performed. The mean voltage at each electrode for each stimulus was computed across the four repetitions. Voltages from different electrodes were concatenated into a row vector and arranged in a neural data matrix with each row containing the vector of voltages for one stimulus sampled from multiple electrodes over time. We rejected columns and then rows where the marginal mean was more than five standard deviations from the grand mean to censor extreme outliers.

### Model fitting and evaluation

RSL decoding models regularized with either grOWL or the L1-norm were fit and analyzed in MATLAB v9.5 (MathWorks, 2018) using the Whole-brain Imaging with Sparse Correlations (WISC) toolbox (Cox, 2021), which implements the approach as a multitask variant of group ordered weighted L1-regularized regression (grOWL; Oswal et al., 2016; Oswal & Nowak, 2018).

Fitting a decoding model regularized with grOWL requires two hyper-parameters that govern pressured the strength with which the decoding matrix is to be row-sparse (λ) with similar weights across correlated neural features (ω). We chose values for λ and ω via ten-fold nested cross-validation for each model fit. The 100 stimuli were randomly divided into ten subsets, each containing five animate and five inanimate items. On each of the ten folds, one of the ten subsets was held out (outer-loop holdout). The remaining nine sets were used to search for good hyperparameter values, involving a second “inner” cross-validation loop.

On each inner-loop fold, one of the nine remaining sets was held out and a decoding model was fit to the remaining eight sets using a specified pair of hyper-parameter values. The fitted model’s prediction error, defined as the Frobenius norm of true versus predicted semantic coordinates, was evaluated on the inner-loop holdout set. This procedure was repeated once for each of the nine inner-loop holdouts before averaging the prediction error across folds to estimate the decoder’s performance with that hyperparameter configuration. Many different hyperparameter configurations were evaluated in the inner loop using the Hyperband procedure (Li et al., 2018). The best-performing configuration was then used to fit a model to all examples not in the outer-loop holdout set. This tuned model was used to predict semantic coordinates for the outer-loop holdout set, thus completing one iteration of outer-loop cross-validation. The whole procedure was then repeated for the remaining nine outer-loop holdout sets. Across folds, the procedure generated out-of-sample predicted coordinates for all 100 items. These final predicted coordinates, computed separately for each time-window in each participant, were the primary data evaluated in the results.

### Statistical thresholding with permutation testing

Our analyses closely followed the simulations: correlations between predicted and true coordinates on each dimension were computed for all 100 stimuli, just the 50 animate items, or just the 50 inanimate items. Assessing model accuracy via correlation standardizes means and variances of the target and predicted vectors to focus on just their covariance. This is especially critical when evaluating animate and inanimate items separately. However, cross-validated correlation has a negative bias when evaluating held-out items (see Zhou et al., 2017 and Appendix C); a null hypothesis of zero with a t-distributed sampling distribution cannot be assumed. Thus, we determined statistical reliability at the group level by constructing an empirical null distribution via a permutation procedure described by Stelzer et al. (2013). The analysis described above was repeated 100 times per participant, each time with a different random permutation applied to the rows of the target matrix. This yielded 100 correlation values for each patient, representing expected values from our workflow when no reliable relationship exists between neural data and target matrix (due to the permutation procedure). Then, a group-level empirical null distribution was estimated by randomly sampling one of the 100 performance metrics from each participant-level distribution and computing a permuted-group-average 10,000 times. This provided sufficient resolution at the group level at a fraction of the computational cost of fitting models to 10,000 permutations of the target matrix per participant for each test (i.e., each window, for each subset of items, in all analyses).

If *m* is the number of values in the permutation distribution and *b* is the number of values in the distribution larger than the true correlation value, then the one-tailed p-value can be computed (Edgington & Onghena, 2007; Phipson & Smyth, 2010) as:

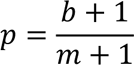

Statistical significance was defined with respect to α < .05 after adjusting the p-values to control the false discovery rate (FDR).

Finally, to understand the effects of regularization with structured sparsity (grOWL) vs. more standard techniques, we compared models fit with grOWL regularization to those fit with L1 (LASSO) regularization.

## Results

### Analysis 1: Full window decoding with grOWL and LASSO

Figure 4a shows results from decoding the full 1000-ms time-window. For the first component, models fit with both grOWL and LASSO reliably decoded similarity structure across all items and for animate and inanimate subsets considered separately. Whereas this was the only structure discovered with LASSO regularization, grOWL regularization additionally showed reliable decoding of the second component across all items and within animates only. Thus, RSL with grOWL regularization revealed graded, multidimensional semantic structure in ECoG signals recorded from ventral ATL, while the contrasting pattern for LASSO suggests that additional constraints from grOWL aided in the discovery of multidimensional semantic structure.

**Figure 4.**
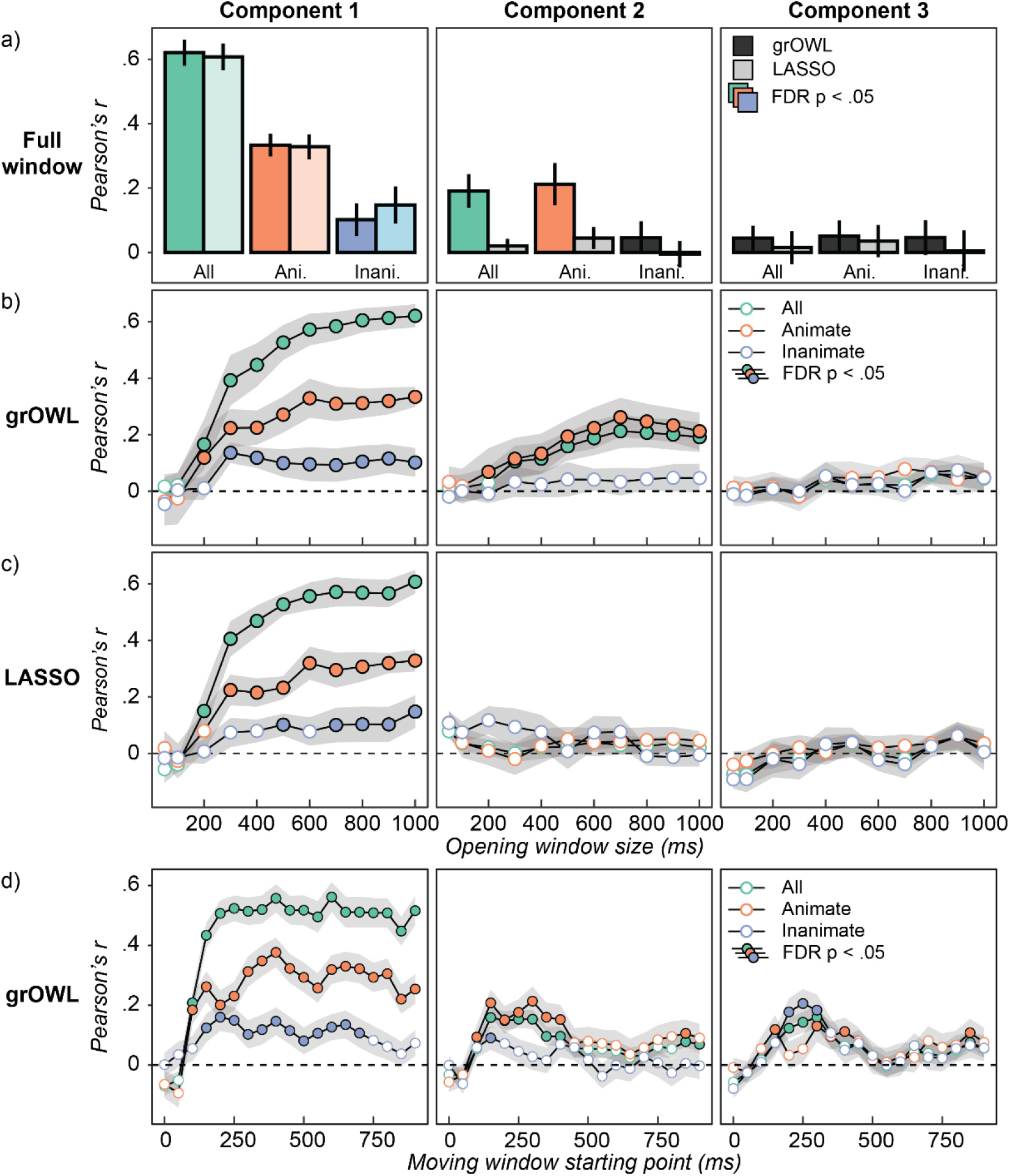
ECoG decoding results. Correlation between true and predicted coordinates along each latent dimension (columns) for models fit to all items (green), animate items only (orange), or inanimate items only (blue), and regularized with grOWL or LASSO. Error bars reflect standard error of the mean over participants. Each value is centered on the mean of its corresponding permutation distribution. For uncentered values, see Figure S4. Colored bars / filled circles indicate reliable decoding with FDR-corrected p < .05 for all points in a panel. a) grOWL (darker bars) and LASSO (lighter bars) model performance when trained on the full 1000-ms trial epoch. b) grOWL model performance within opening windows. c) LASSO model performance within opening windows. d) Analogous to (b) except that models are fit and evaluated within a 100ms moving window instead of the opening window.

### Analysis 2: Opening window

To see how this structure emerges in the ECoG signal over time, we conducted an “opening window” analysis in which the same procedure was applied to an increasingly wide aperture of data, beginning with just the first 50ms post-stimulus, extending to 100ms, then growing by 100ms up to 1000ms. The opening window analysis evaluates when enough information has entered the spatiotemporal feature space to support decoding. The goal is not to localize a representation in time, but to identify when reliable decoding is first possible and when performance stops improving.

The results show reliable decoding of between-domain and within-animate structure along the first component by 200ms, followed by within-inanimate structure by 300ms (Figure 4b). Decoding accuracy for superordinate and animate-subordinate structure continually improved with wider windows along component 1, but not inanimate-subordinate structure. Along the second component, reliable decoding was observed after 300ms for between-domain structure and somewhat earlier (200ms) for within-animate structure. Within-inanimate structure could not be reliably decoded along the second component at any window-size, nor could variation along the third component of the target matrix. Models fit with LASSO (Figure 4c) also reliably decoded both within- and across-domain structure along the first component, beginning at 200ms for cross-domain structure, 300ms for a within-animate structure, and 500ms for within-inanimate structure, but as with the full window, did not reliably decode the second (or third) component for any window size.

### Analysis 3: Moving window

The opening window analysis indicates the latency with which the neural signal contains sufficient information for reliable decoding, but since each successive window contains all prior time points, it does not indicate whether/how the neural encoding of semantic information changes over time. Additionally, since larger windows contain more neural features, they afford a greater possibility of over-fitting training data. Consequently, the opening-window approach may fail to detect semantic information encoded only within a limited time-window. For these reasons, we fit models using the grOWL regularizer on a 100ms *moving* window, beginning at 0ms from stimulus onset and advancing in 50ms increments. In this analysis, each window is the same size and thus contains the same amount of neural data. Otherwise, the analysis was identical to the opening-window variant.

Results are shown in Figure 4d. For component 1, reliable decoding was observed across domains and within each domain between 150 – 700ms post stimulus onset. Reliable decoding on components 2 and 3 was observed within a more limited time range. For component 2 (which best-separates the animate items), cross-domain and within-animate structure was reliably and equally-well decoded for windows beginning at 150 – 400ms. For component 3 (which best separates inanimate items), cross-domain and within-inanimate structure was reliably and equally-well decoded for windows beginning at 200 – 300ms. Together the analyses suggest that, from around 200 – 400ms post-stimulus, neural states measured by ECoG express both within- and between-domain semantic structure, for both animate and inanimate items, across all three components of the target matrix. That is, they express a graded multidimensional and domain-general semantic space.

### Analysis 4: Moving-window reconstruction of full target similarity matrix

The preceding analyses consider decoding of each matrix component separately, allowing us to draw conclusions about which aspects of semantic structure are represented in the neural signal at which time points. The independent consideration of different dimensions, however, makes it difficult to compare results of RSL to RSA, since RSA considers correlations between full pairwise similarity matrices (the NSM and the RSM). To facilitate the comparison, we used the decoding models fit in Analysis 3 to construct a *predicted semantic similarity matrix* at each time-window. For each fitted model, we predicted coordinates of the corresponding held-out items along all three target matrix components, agglomerating these predictions across hold-out sets to create a matrix of predicted coordinates for all items. Recall that the target coordinates are the first three singular vectors of the original semantic similarity matrix, weighted by the *square root* of the corresponding singular values (Figure 3). Thus, to reconstruct predicted pairwise distances in the original target matrix, we need only take the product of the predicted-coordinates matrix with its transpose.

Each row of the resulting matrix contains predicted semantic similarities between an item and all other items. We then compared these predicted similarities to a target similarity matrix constructed directly from the three components of the original matrix that account for 90% of its variance. That is, for each item in the dataset, we computed the correlation between the predicted and true similarities to (a) all other items, (b) other items in the same domain, and (c) other items in the contrasting domain. For each metric, we then averaged the correlations across (1) all items, (2) just the animate items or (3) just the inanimate items. These conditions thus allowed us to assess how well the decoders model semantic similarities both within and across each domain. The full procedure was carried out independently for each participant at each time-window.

Correlation coefficients in each condition were then averaged across participants at each time-window.

The results are shown in Figure 5. Filled circles indicate where predicted/true correlations are reliably non-zero relative to a permutation-based null distribution with FDR of p < 0.05 (Stelzer et al., 2013). Reliable correlations were observed from 100 – 150ms post-stimulus onward, whether computed across all items, within domain only, or between-domain only, and considering the full complement of items (left panel), animate items only (middle), or inanimates only (right).

**Figure 5.**
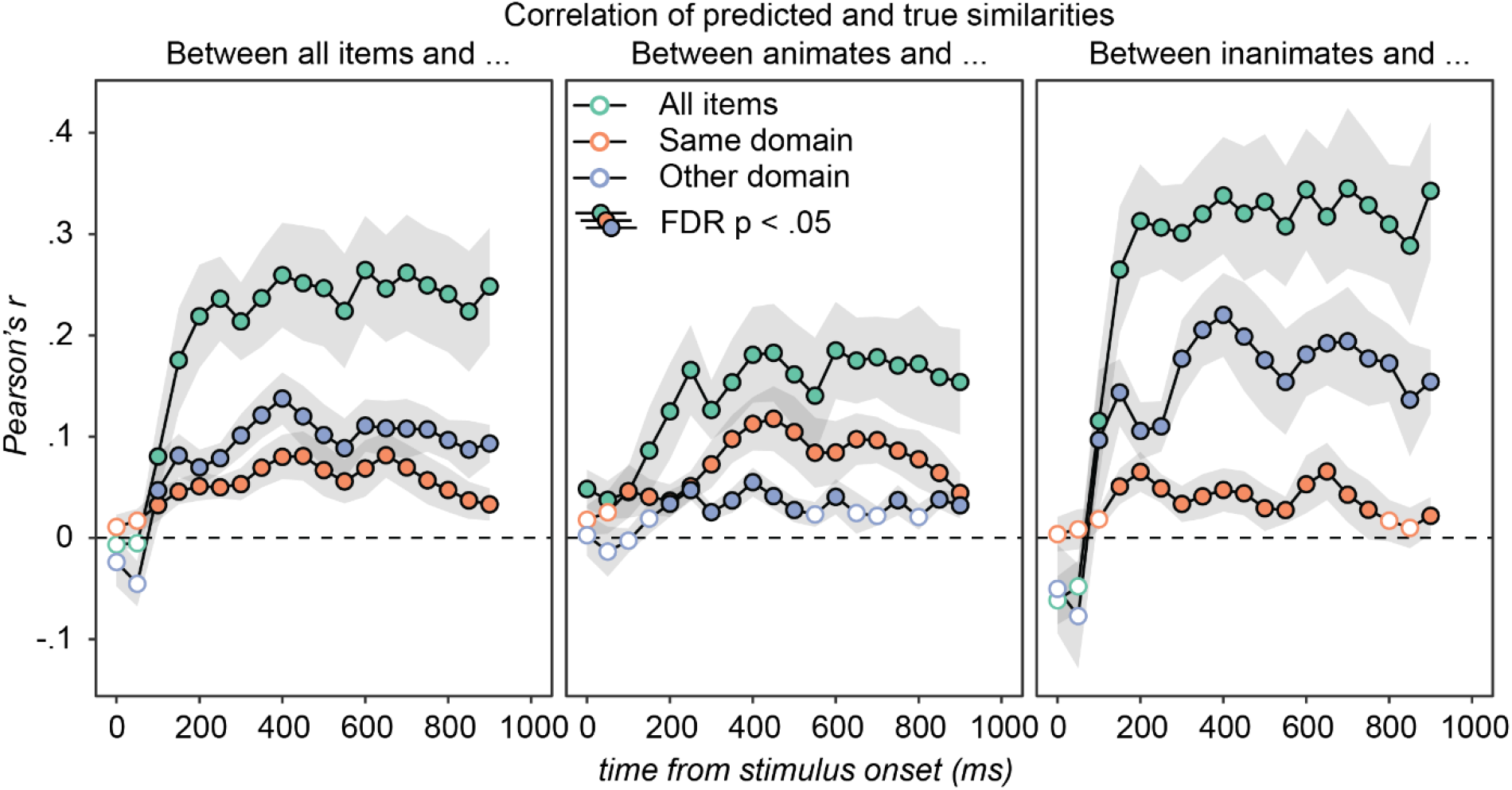
Correlation between predicted semantic similarities and the best-possible similarities reconstructed from the first three singular vectors/values of the true matrix. Error bars reflect standard error of the mean over participants. Decoding models fit with RSL reliably predict semantic similarities both within (orange dots) and between (blue dots) conceptual domains, for both animate (middle panel) and inanimate (right panel) stimuli, from about 100 – 200ms post-stimulus onward.

For comparison, we replicated the RSA analysis of Y. Chen et al (2016) on the same data used for RSL and extended it to examine structure within animate and inanimate item subsets. When conducting RSA over all items, NSMs were correlated with both a multidimensional target RSM and a unidimensional categorical (i.e., binary) target RSM. Figure 6 (left) plots the average Spearman’s rho over participants within the same ROI reported by Y. Chen et al (2016) for each temporal window and each target RSM in light and dark grey. The correlations with each target RSM are very similar, consistent with the original finding that RSA did not provide clear evidence about the kind of semantic information present in the vATL. The right panel shows results of RSA applied to animate (light grey) and inanimate (dark grey) item subsets. These correlations were indistinguishable from zero in every window.

**Figure 6.**
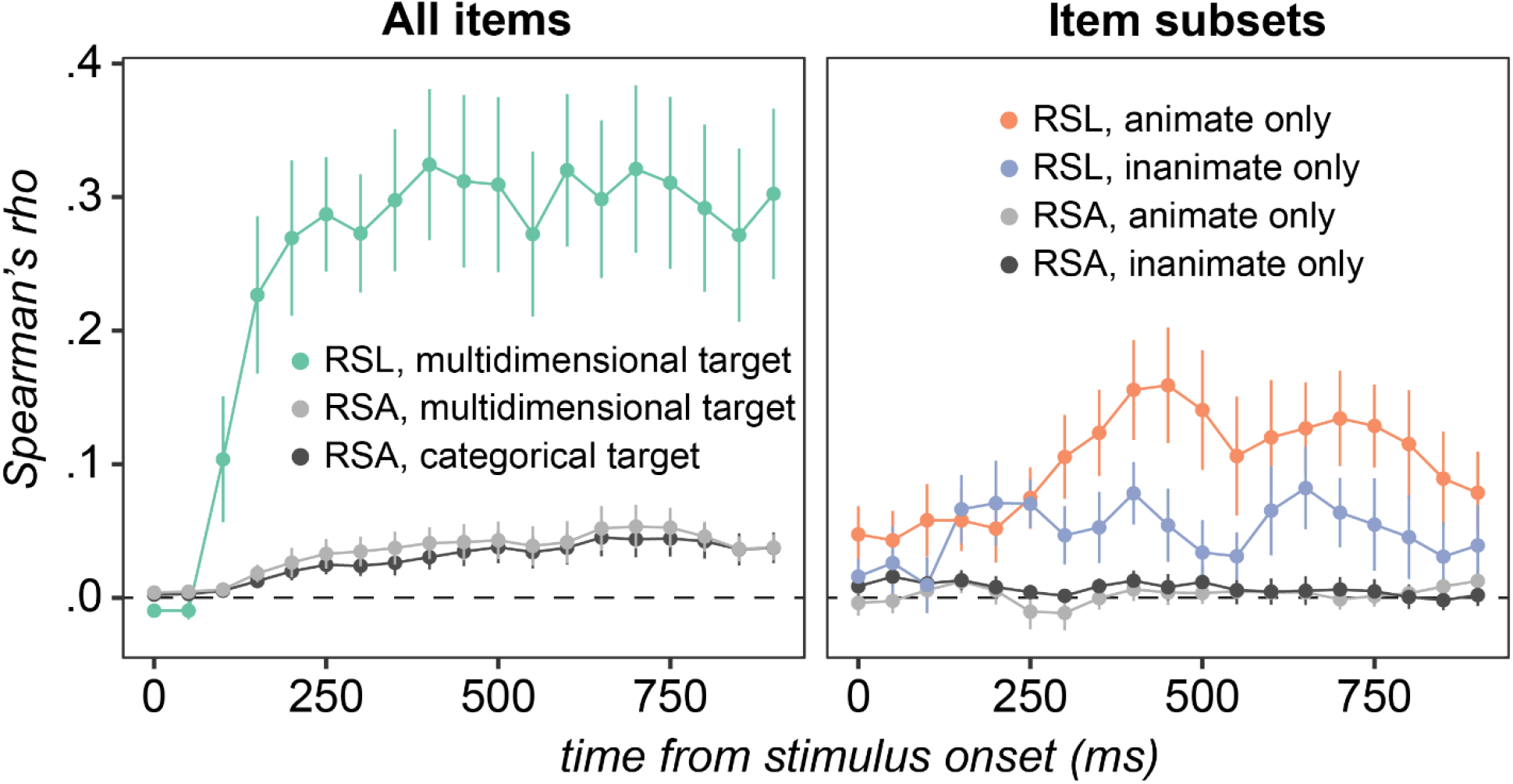
Comparing RSA and RSL on the same data. All correlations (Spearman’s rho) are averages over the 10 participants and error bars depict standard error. The left panel shows the correlation between the full predicted similarity matrix obtained using RSL and the multidimensional target RSM (green), as well as results obtained by replicating the RSA conducted by Y. Chen et al (2016) on the same data used for RSL using the multidimensional (light grey) or a categorical (dark grey) target RSM. Differences between categorical and multidimensional RSA are insignificant, while differences between RSL and RSA are large and significant from 150ms onward. The RSA results in the right panel are analogous to the left panel except that subsets of items are studied separately. RSA does not detect within-category structure while RSL does.

Colored lines in the two panels show the correlations between the RSL predicted similarity matrix and the target matrix across all items (left panel) or within animates (right panel orange) and within inanimates (right panel blue). While reliably positive correlations over the full set of items were observed with both RSA and RSL from 150ms onward (one-sample two-tailed t-tests, μ_0_ = 0, α = .05, all t(9)>2.75), the correlation coefficients obtained by RSL were significantly larger in all windows with reliable effects (paired two-tailed t-tests, α = .05, all t(9)>3.5). However, this is not a difference that requires a statistical analysis to appreciate: over windows with reliably positive correlations, the correlation coefficient obtained with RSL was between 6 to 13 times larger. A 10-fold increase in Spearman’s rho is a 100-fold increase in the proportion of variance explained. RSL produces a qualitatively different result from RSA, in terms of its sensitivity to detecting structure and its ability to determine what structure is present in neural signal.

## Discussion

Replicating Y. Chen et al. (2016), if one only relied on RSA analysis then it would be easy to conclude that ECoG activity in vATL only encodes a discrete, binary distinction between animate vs. inanimate items, with no information about semantic structure within either domain considered independently. In stark contrast, the RSL analysis shows that ECoG signals measured in human vATL encode information about semantic similarity structure that is multidimensional (reliable decoding along three orthogonal components of the target matrix), graded (reliable decoding of varying degrees of similarity both within and between domains), and domain-general (reliable decoding of within-domain similarities for both animate and inanimate items). These properties are consistent with the predictions of the “hub and spokes” theory of semantic representation in the brain, which proposes that neural activity in vATL encodes a graded, multidimensional, and domain-general semantic vector space. It is also consistent with the converging sources of evidence that gave rise to that view, including patient studies (Patterson et al., 2006), neural stimulation (Pobric et al., 2007), and computational modeling (Rogers et al., 2004). In this sense, the RSL results resolves a seeming discrepancy between RSA findings and the broader literature.

## General Discussion

We introduced this paper with a puzzle: neuropsychology, clinical neurophysiology, TMS, and computational modelling all suggest that the vATLs encode a semantic vector space of graded, multidimensional and domain-general conceptual similarity structure, but direct tests of this hypothesis using representational similarity analysis have often yielded null results in vATL and positive results in brain areas not otherwise thought to encode semantic representations. In simulation we showed that counter-intuitive limitations of RSA can obscure inferences about neurocognitive representation. When used to decode real semantic structure as measured by feature norms, RSA can produce positive results even if the underlying neural code is discrete and unidimensional, or null results even if the neural code does capture latent structure in the target matrix beyond the first component. Because RSA relies on an inherently unidimensional measure of association (i.e., correlation), it cannot reveal whether neural signals encode multidimensional structure. Because the technique does not fit any parameters to data, it requires the researcher to select features a priori when constructing the NSM (for instance, via ROI or searchlight analyses; see Frisby et al., 2023), and cannot learn to ignore irrelevant features amongst those selected. The simulations suggest that these characteristics of RSA may have contributed to the puzzling state of the literature—for instance, by yielding results suggesting that vATL only coarsely discriminates living from nonliving things (Clark & Tyler, 2014; Chen et al., 2015).

RSL addresses these limitations by using regression to predict coordinates of stimuli along the latent orthogonal dimensions of the RSM. In simulation, we showed how this approach can uncover multidimensional representational structure (by showing reliable conjoint decoding of two or more orthogonal dimensions of a target matrix) as well as graded structure (by showing reliable decoding both within and between semantic domains). The approach can also “select out” signal-carrying features from amongst those included as predictors in the model, and so can be applied to all potentially signal-carrying neural features at once, without requiring the theorist to pre-select an ROI or to look only within small, independent searchlights (see Cox & Rogers, 2021; Frisby et al., 2023).

Applying the approach to large neural datasets requires model regularization. We illustrated how hypothesized patterns of structured sparsity in the neural signal can constrain model fit via grOWL regularization and applied this approach to discover semantic structure in an ECoG dataset where prior work using RSA found only a binary animacy code (Chen et al., 2015). We replicated this analysis using the same RSA approach, and further showed that RSA yields null results when assessing whether the ECoG signals express within-domain semantic structure. In contrast, RSL with grOWL regularization uncovered a graded and multidimensional semantic space capturing similarities within and between both animate and inanimate domains—consistent with conclusions drawn about the nature of semantic representations in the ATL hub from other cognitive and clinical neuroscience sources. These results thus suggest that discrepancies in the literature between studies employing RSA versus other sources of evidence may reflect limitations of the RSA approach as typically practiced.

### Validity of grOWL assumptions about neural signal

The RSL models we have deployed were fit with a regularization function designed to promote discovery of a *row-sparse* decoding matrix (the grOWL loss). We hypothesize that such structure reflects three characteristics of neuro-semantic representation specifically, and neuro-cognitive codes generally. First, the signal is likely to be sparse: of all neural measurements taken in a given experiment, only a relatively small proportion are likely to encode the target information of interest. The sparsity assumption serves a useful role in decoding because it pressures many coefficients in the regression model to zero, indicating that the corresponding features are not useful in decoding the target information. In this sense, sparsity automatically serves the function of feature-selection that, in RSA, must be handled a-priori based on an ROI, searchlight, or other method (Cox & Rogers, 2021). We hypothesize neural codes to be relatively sparse in general simply because brains support many different cognitive, perceptual, motor, language, and affective functions—consequently the likelihood that a given neural population is important for the specific function targeted by the investigator is relatively small.

Second, we hypothesize that the signal is *redundant*: it is unlikely that a given target structure is only encoded by a single voxel, or a single timepoint at a single electrode, etc. Any cognitive construct of interest—feature, category, or dimension in a representational space—is unlikely to be encoded by the activation of a single local neural population, such as a single voxel. More likely, such information involves multiple neural populations, in which case those populations that *do* encode target information will exhibit some degree of correlation with one another. This assumption is captured by the grOWL loss because it encourages solutions in which intercorrelated sets of neural features that help to predict target structure receive similar, non-zero coefficients in the decoder.

Finally, we hypothesize that the neural code is unlikely to be axis-aligned with the dimensions of the target representation space. That is, a given neural feature is unlikely to encode variation along just one dimension of a multidimensional target space without also encoding some information about variance along other dimensions. In grOWL, this hypothesis is expressed as a preference for learning a row-sparse decoding matrix— coefficients on a given neural feature should either be all zero (the feature is unimportant) or all non-zero (the feature explains some variance along each dimension). The reason is simply that, for any target vector space, there exists only a small and finite number of axis-aligned encodings, but an infinite number of non-axis-aligned encodings. Aligned and unaligned encodings express the same information about similarities amongst objects of representation, so, absent some explicit pressure for brains to learn axis-aligned representations, it is unlikely that an axis-aligned encoding will occur by chance. Additionally, ECoG electrodes are influenced by voltages generated by a mix of individual local neurons—so even if individual neurons are selectively tuned to axis-aligned dimensions, the net responses recorded at the electrode is likely to reflect a blend of these dimensions.

While the grOWL regularizer generally prefers decoding models with these properties, note that the relative strength of these constraints is determined by hyperparameters that can be tuned via cross-validation to fit the data. If the best solution is not particularly sparse, the tuning process will select a small weight for the sparsity term in the optimization, leading to a solution in which many features are chosen. If the signal-carrying features are not particularly correlated, the tuning process can select hyperparameters that relax the grouping of features into sets that share the same weights. Thus, grOWL regularization is quite flexible with respect to how rigidly the various constraints are enforced.

The current study suggests that the grOWL assumptions are useful for understanding information encoded in ECoG voltages: when decoding the full time-window, regularization with grOWL revealed multidimensional similarity structure, whereas decoding with the sparsity assumption only (LASSO) found only unidimensional structure. We emphasize, however, that the RSL framework can be deployed with any form of model regularization. Alternative hypotheses about the likely structure of neural encodings can be formulated as different regularization costs, and the decoding success of models fit with different regularizations can then be compared to empirically evaluate the different assumptions. Prior computational work has contrasted grOWL regularization to other approaches (see Oswal et al., 2016 for comparisons to some other related methods); we hope the current results with grOWL will inspire other scientists to experiment with alternative losses to better understand the nature of neuro-cognitive codes.

### Implications for alternative theories of ATL function

The current results challenge an alternative proposal about the role of ATL in semantic cognition, namely that they support knowledge of some conceptual categories and not others (Mahon & Caramazza, 2011; Malone et al., 2016; Simmons et al., 2009). Such a view is consistent with prior imaging work (e.g., Anzellotti et al., 2011) including the multivariate decoding studies cited earlier (Y. Chen et al., 2016; Clarke & Tyler, 2014), suggesting that ATL activations differentiate animates from inanimates but do not otherwise encode or differentiate finer categories. This alternative perspective has been difficult to reconcile with neuropsychological evidence showing that anterior temporal atrophy and hypometabolism in semantic dementia degrades knowledge of animates and inanimates equally (Lambon Ralph et al., 2007; Noppeney et al., 2007), and with TMS and direct grid stimulation evidence showing that stimulation of both left and right ATL reliably slows semantic processing for both animates and artefacts (Pobric et al., 2010; Shimotake et al., 2015). It also struggles to explain the sensitivity of ATL-related semantic impairment to continuous and graded semantic structure of test stimuli for both animates and inanimates (Patterson et al., 2006; Rogers et al., 2006). The current results suggest that prior work may have failed to discover graded cross-domain semantic structure in ATL responses, not because such structure is absent from the measured responses, but because limitations in both fMRI and analytic techniques make such structure difficult to detect.

It is worth noting that intra-domain similarities were more robustly decoded for animate compared to inanimate items. Considering each dimension separately in the opening window analysis, within-inanimate decoding was only reliable for the first latent dimension. In the moving window analysis, where models were trained on 100ms increments of data without the full history of activation from stimulus onset, within-inanimate decoding was reliable for components 1 and 3, but transiently. These are the components that best differentiate inanimates in the target matrix (see Figure 1). When predicting pairwise distances within moving windows, correlations with true similarities were reliable but smallest for within-inanimate structure (i.e., the “other domain” correlations in the middle panel of Figure 5, and “same domain” correlations in the rightmost panel).

We attribute this general pattern to differences, not in the neural code itself, but in the target similarity structure of animate and inanimate concepts. Semantic subcategories of inanimate objects are only weakly differentiated, not just in the current dataset, but in norming studies more generally and in other approaches to characterizing semantic structure (e.g., Devlin et al., 1998; Garrard et al., 2001; McRae et al., 1997; Tyler et al., 2000). Indeed, this difference in statistical structure is precisely what leads, under some theories, to apparent category-specific patterns of semantic impairment (Devlin et al., 2002; Lambon Ralph et al., 2007) and functional activation (Rogers et al., 2006). The differences can be seen in Figure 1, where animate items fall into somewhat well-differentiated subcategories in components 1 and 2 while inanimates are uniformly distributed with poorly differentiated subclusters even along component 3. Accordingly, pairwise similarities reconstructed from the first three singular vectors of the full semantic matrix are more accurate for animate than inanimate items (see Appendix D). Thus, the structure of the semantic matrix itself *requires* that decoding be worse within inanimates than within animates, precisely because inanimates are less well-structured. Yet despite this intrinsic disadvantage, within-inanimate similarities were still reliably decoded along multiple components of the embedding and in the reconstructed similarity matrix, illustrating that neural signal in vATL does express intra-domain semantic structure even for inanimate stimuli.

In summary, the results suggest the vATLs encode a multidimensional representation space that captures the conceptual similarities existing amongst a variety of different concepts including both animate and inanimate items (Lambon Ralph et al., 2007; Patterson et al., 2007; Rogers & McClelland, 2004). Better decoding within animates may reflect the fact that animate subcategories are better-differentiated, so that intra-domain similarities are better-approximated by a low-rank decomposition.

We also note that the current results pertain only to the nature of the semantic information encoded within the span of vATL where electrodes were situated in our patient sample. They do not bear on claims of possible category-specific representation in other parts of cortex—for instance, that left infero-parietal cortex and posterior medial temporal gyrus play a special role in knowledge about tools (Q. Chen et al., 2016; Garcea et al., 2018; Ishibashi et al., 2016; Kalénine & Buxbaum, 2016) or that earlier visual areas are dedicated to representing different object categories (Cichy et al., 2014; Connolly et al., 2012; Downing et al., 2001; Kanwisher et al., 1997; Kriegeskorte, Mur, Ruff, et al., 2008; Mahon et al., 2009; Sha et al., 2015). Indeed, prior work from (Chen et al., 2017) showed how domain-general semantic representations can arise in vATL even as graded category-specificity emerges in other parts of the cortical semantic network, based on empirically-measured patterns of white-matter connectivity across core areas. Evaluating claims of category-specificity in future work may benefit from adopting the RSL approach developed here.

### Implications for the broader literature

The characteristics of RSA we have identified carry additional implications for interpretation of the broader literature. A recent review (Frisby et al., 2023) identified 24 studies that have applied RSA to the discovery of semantic representations in the brain, with positive results observed across multiple cortical regions including posterior temporal cortex (Connolly et al., 2012), angular gyrus (Fairhall & Caramazza, 2013; Fernandino et al., 2022), left perisylvian cortex (Devereux et al., 2013), posterior cingulate (Fairhall & Caramazza, 2013), and prefrontal cortex (Carota et al., 2017). Simulation 1 suggests, however, that RSA will yield a positive result for any property correlated with the animate/inanimate distinction, including even discrete binary properties. Many features are confounded with animacy: inanimate items tend to be more familiar, less visually complex, more associated with action plans, less associated with motion, more likely to have lines and corners, less predictable from color or texture, etc. (see Chen & Rogers, 2014 for a review). Most RSA papers do not report the magnitude of correlation between RSM and NSM where results are significant, instead focusing on whether the estimated correlation coefficient is reliably non-zero across participants. As shown by the prior RSA analysis of the same ECoG data explored here (Y. Chen et al., 2016; gray line in Figure 5), this can happen even when mean correlations are very small. Moreover, several studies use target RSMs with only a small number of rows/columns—sometimes as few as five (Connolly et al., 2012; Fairhall & Caramazza, 2013), meaning that correlations are computed across just 10 cells of the matrix (i.e., the lower triangle of a 5×5 RSM). Such small numbers increase the likelihood that a small-but-non-zero correlation is driven by some arbitrary property of the chosen stimuli or categories. Together these observations raise the possibility that the literature contains misleading positive results—brain areas whose responses encode unidimensional characteristics weakly confounded with animacy, rather than multidimensional semantic structure. Testing this possibility for various brain areas hypothesized to encode semantic structure will require analyses like those we have developed here.

### Conclusion

In cognitive science, semantic representations are often construed as vector spaces that encoded graded, multidimensional similarity structure amongst the concepts experienced in our verbal and nonverbal world. Through application of a new technique for mapping representational similarity in neural activity, we have shown that neural signals in vATL encode such a space. In so doing, we have identified some limitations of representational similarity analysis, a widespread technique commonly thought to reveal graded and multidimensional representational structure. The work resolves an important discrepancy between behavioral and neuroimaging results in prior work and suggests a new approach to discovering representational structure in neural data more generally.

## Author Contributions

Cox conducted all analysis and drafted the manuscript. Rogers and Lambon Ralph contributed to all aspects of the study design, analysis, and manuscript preparation. Shimotake, Kikuchi, Kunieda, Miyamoto, Takahasi, Ikeda and Matsumoto were responsible for ECoG data collection and clinical assessment.

## Data and materials availability

RSL models were fit using the WISC MVPA toolbox (https://github.com/crcox/WISC_MVPA). Analysis of RSL models was conducted using custom MATLAB code (https://github.com/crcox/RSL_correlation_analysis), and the data and script necessary to replicate them is available via the Open Science Framework (https://osf.io/axseq/). R scripts for generating figures based on the correlation analyses are on GitHub (https://github.com/crcox/ecog-rsl-figures).

## Appendices

Appendix A. Categorized experimental stimuli

Appendix B. Formal description of the RSL approach

Appendix C. RSL and negative correlations Anatomical distribution of decoding model weights across vATL

Appendix D. Reconstructing item similarity from the SVD coordinates

Appendix E. Correlation between predicted and true semantic distances

Appendix F. Temporal Generalization profiles of models fit within moving windows

Appendix G. Anatomical distribution of decoding model weights across vATL

Appendix H. Predicted embeddings

## Appendix A: Categorized experimental stimuli

**Table S1.** Categorized experimental stimuli.

| **Domain** | **Category** | **Words** |
| --- | --- | --- |
| *Animate* | *Bird* | duck, eagle, ostrich, owl, peacock, penguin, rooster, swan |
|  | *Bug* | ant, bee, butterfly, caterpillar, fly, ladybird, snail, spider |
|  | *Land* | bat, bear, camel, cat, cow, deer, dog, dragon, elephant, fox, frog, giraffe, goat, gorilla, horse, kangaroo, lion, monkey, mouse, pig, rabbit, raccoon, rhinoceros, sheep, snake, squirrel, tiger, tortoise, zebra |
|  | *Water* | crab, fish, lobster, seahorse, whale |
| *Inanimate* | *Clothes* | dress, glove, mitten, skirt |
|  | *Music* | bell, cymbals, drum, flute, guitar, harp, piano, trumpet, violin, whistle |
|  | *Object* | anchor, barrel, basket, cake, cannon, chain, church, cigar, clock, crown, fence, gun, hammer, iron, kite, ladder, nut, pram, shell, slide, snowman, suitcase, swing, toaster, vase, web, wheel, windmill, yoyo |
|  | *Vehicle* | aeroplane, helicopter, motorcycle, ski, sled, tractor, train |

## Appendix B: Formal description of the RSL approach

Let ***S*** be a symmetric *n* × *n* target matrix of approximate rank *r* describing pairwise cognitive similarities amongst *n* items, which is decomposed via singular-value decomposition (SVD) into an *n* × *r* matrix ***U*** of *r* orthonormal singular vectors and an *r* × *r* diagonal matrix ***D*** containing the corresponding *r* singular values (**Figure 2A**). The matrix product 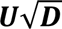 then contains root-weighted singular vectors that can be viewed as encoding the coordinates of each stimulus within an *r*-dimensional latent semantic space, while the product of this matrix with its transpose provides a rank *r* approximation of the original similarity matrix. That is:

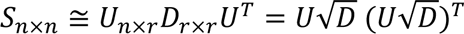

Let ***X***_*n*×*m*_ be a data matrix recording neural responses to each of *n* stimuli over *m* neural features (e.g., voxels in fMRI, electrode voltages at each timepoint in ECoG, etc.). To assess whether some weighted combination of these features jointly encode the target similarities, we seek a *decoding matrix* **β**_*m*×*r*_ such that the matrix product ***X*β** produces *r* coordinates for each item, deviating as little as possible from those in target matrix 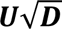. This search can be formulated as the convex optimization:

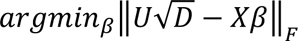

That is, we seek **β** values that minimize the Froebenius norm (i.e., sum of squared differences) between the true root-weighted singular vectors 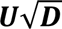 and the predicted values ***X*β**. The optimization essentially computes a simultaneous regression predicting all *r* coordinates for each stimulus from weighted combinations of the neural responses they evoke. Each column of the resulting decoding matrix contains one coefficient for each neural feature and predicts an item’s coordinate along one component of the target matrix 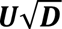 (ie, one dimension of the semantic space).

As noted earlier, there are typically many more neural measurements (predictors) than stimuli (i.e., target data points)—that is, *m* ≫ *n*. Thus, the optimization just defined is under-constrained and must be regularized with an additional loss *H*, itself a function of the coefficient matrix **β**:

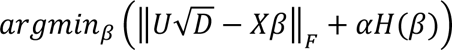

…which is also convex so that a weighted sum of this function and the original reconstruction loss together yield a single unique solution that can be discovered via gradient descent. The constant α is a free weighting parameter governing the importance of the regularization loss relative to the model error.

For RSL we wish to use a regularization function that promotes solutions respecting the sparsity, redundancy, and spanning assumptions described earlier. Oswal et al. (2016) formulated a regularizer that accomplishes these objectives by constraining the structure of the decoding matrix **β** in a single convex loss called the *group ordered weighted LASSO* or *grOWL.* The approach generalizes both the group LASSO (Yuan & Lin, 2006) and the ordered-weighted LASSO (Figueiredo & Nowak, 2016) to favor solutions that are (1) row-sparse (rows of **β** are either all zero or all non-zero), implementing both the sparsity and spanning constraints, but (2) with subgroups of selected features sharing similar coefficients if their activations covary strongly together (implementing the redundancy constraint). The grOWL loss is specified as:

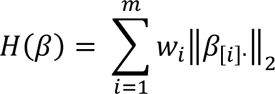

Here *m* is the number of neural features (rows of **β**), **β**_[*i*]_. is the row of **β** with the *i*-th largest 2-norm (i.e., the *i*^th^ longest coefficient vector), and *w*_*i*_ is a vector of non-negative, non-increasing weights (for instance a vector of positive values that decay linearly from an initial constant). Intuitively the loss works as follows. The first element of the weighting vector *w* is stipulated to be as large or larger than all other elements, and this weight is applied to whichever neural feature has the largest coefficients in the decoding matrix (i.e., the 2-norm or length of the corresponding row). Thus, the feature with the largest coefficients exerts the largest regularization cost. The optimization must therefore put the largest weights on the most informative feature to make it “worth” the high regularization cost. If the most informative feature is also correlated with many other features, their total cost can be further minimized by placing similar weights across all such features, since the cost for each such weight declines with the weight sequence in *w*. Thus, grOWL spontaneously clusters correlated signal-carrying neural features and gives these a similar weight, effectively averaging them. Finally, the weighting vector ensures that coefficients on features that do not reduce error beyond those already selected will be pressured to zero, ensuring overall sparsity. Formal analysis of the grOWL regularizer and mathematical proofs of its row-sparse and clustering properties were provided by Oswal et al. (2016).

## Appendix C: RSL and negative correlations

The analysis protocol presented in this paper involves generating out-of-sample predictions with k-fold cross validation and then aggregating those test-set predictions into a single matrix before correlating with the target embedding all in one go. This is in contrast with a more conventional protocol in which the performance metric is computed separately within each fold and then averaged. When the metric is something like mean squared error or classification accuracy, the two protocols are equivalent. With correlation, however, the two protocols yield different values.

Pearson’s r compares variables with zero mean and unit variance. Consequently, when correlations between target and predicted embeddings are computed separately by holdout-set, the scale of the predictions relative to the target is disregarded. By aggregating over test sets and correlating one time, the standardization is done at the level of the full dataset and not for individual test sets. We feel this is a better metric of whether our models can embed all items appropriately within the target similarity space—it matters how items are situated with respect to one another even if they are in different test sets.

When models are fit with cross validation, the targets (and the data) are standardized with respect to the values in the training set. Thus, the model predictions, including test set predictions, are on the scale of the standardized *training set*. This means that, to aggregate over test sets and correlate with the full set of targets, the mean and variance of the target embeddings in the *training set* need to be added back into the predictions for the corresponding *test set*.

Applying this adjustment will introduce negative correlations between target and predicted embeddings for models that have been regularized to have weights that are all zero. This will happen when relationships between the neural data and the target embedding is weak or non-existent, because putting a weight on any neural feature increases the model cost function without reducing the prediction error on the training set. In such cases, the unadjusted model predictions will all be zero, and the adjusted predictions will be the mean of the training set. If the mean of the training set is greater than the grand mean of the full set, then the mean of the test set must be less than the grand mean of the full set. Thus, adjusted and aggregated test set predictions will be negatively correlated with the full target embedding when the model has all zero (or trivially small) weights due to the LASSO or grOWL regularization penalty. These concepts are illustrated in Figure S1 in an imagined case where all-zero models are learned on each of three cross-validation folds.

What we found in practice is that grOWL is “more aggressive” about assigning zero or trivially small weights than LASSO in the earliest and smallest time-windows. We consider this to be desirable behavior because we do not expect much semantic activity in the ATLs at 50ms post stimulus onset. Figure S2 is analogous to Figure 4 in the main paper, except that the correlation values are not centered on the mean of their corresponding permutation distribution. This reveals the negative correlations between grOWL model predictions and the target embedding for small windows where we do not expect semantic activation to be reliably present, and general consistency of performance elsewhere. These negative correlations are an expected artifact of our analysis protocol and do not change the interpretation of our results.

**Figure S1.**
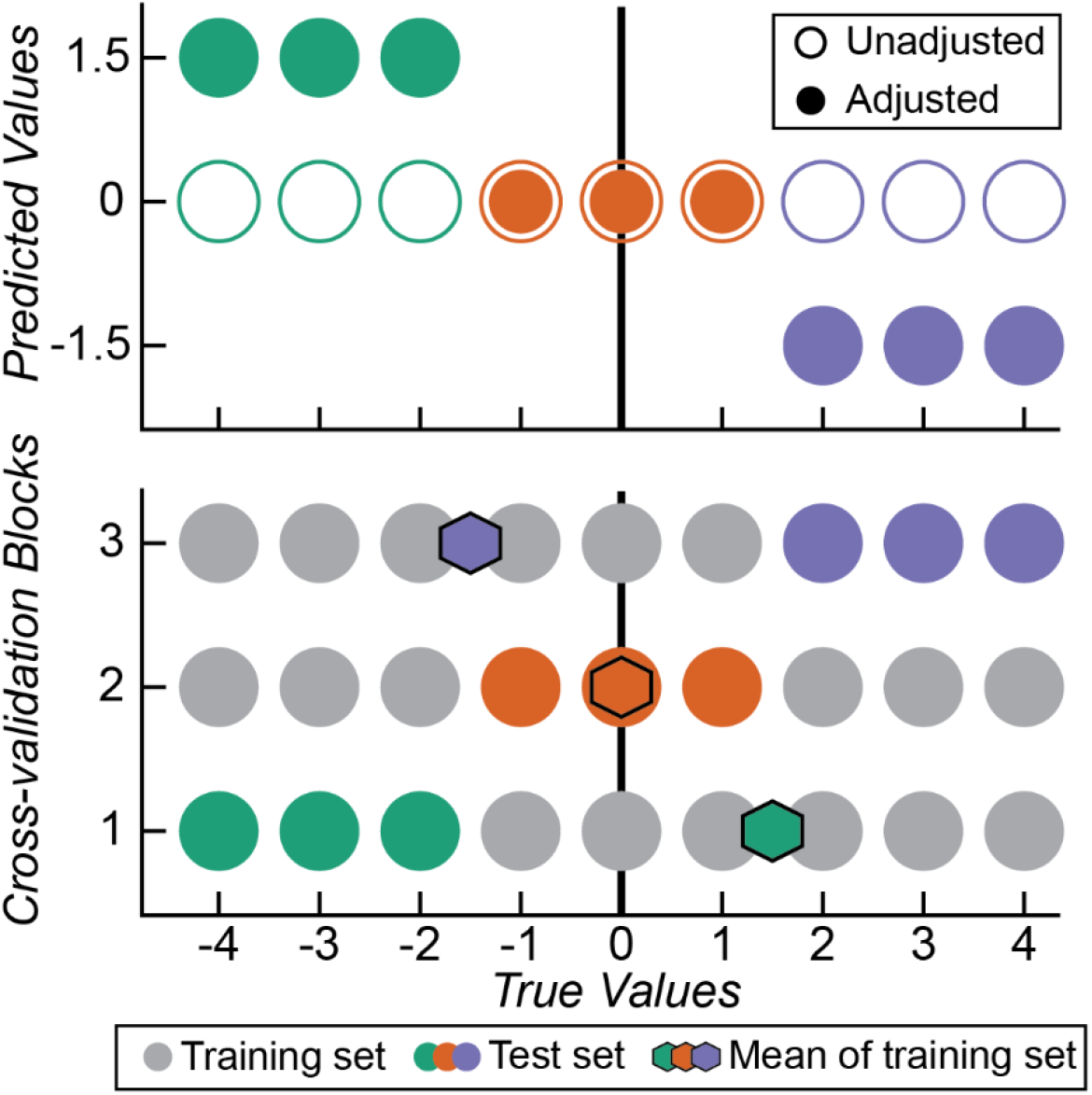
A schematic depiction of the relationship between test sets, training set means, the sample grand mean, and model predictions when the model consists of zero-weights on all neural features. A zero-weight model can occur when model regularization involves the LASSO penalty and the cost of applying a non-zero weight does not outweigh the benefit of reducing prediction error. If the mean of the test set differs from the grand mean, the mean of the test set must also differ from the grand mean. Importantly, if the mean of the test set is less than the grand mean, the mean of the training set must be higher. Unadjusted predictions (i.e., those that ignore the mean of the training set) will be all zero when the model weights are all zero; adjusted predictions are scaled by the training set mean. Leads to systematically negative correlations between true and predicted values for models regularized to have all-zero weights.

**Figure S2.**
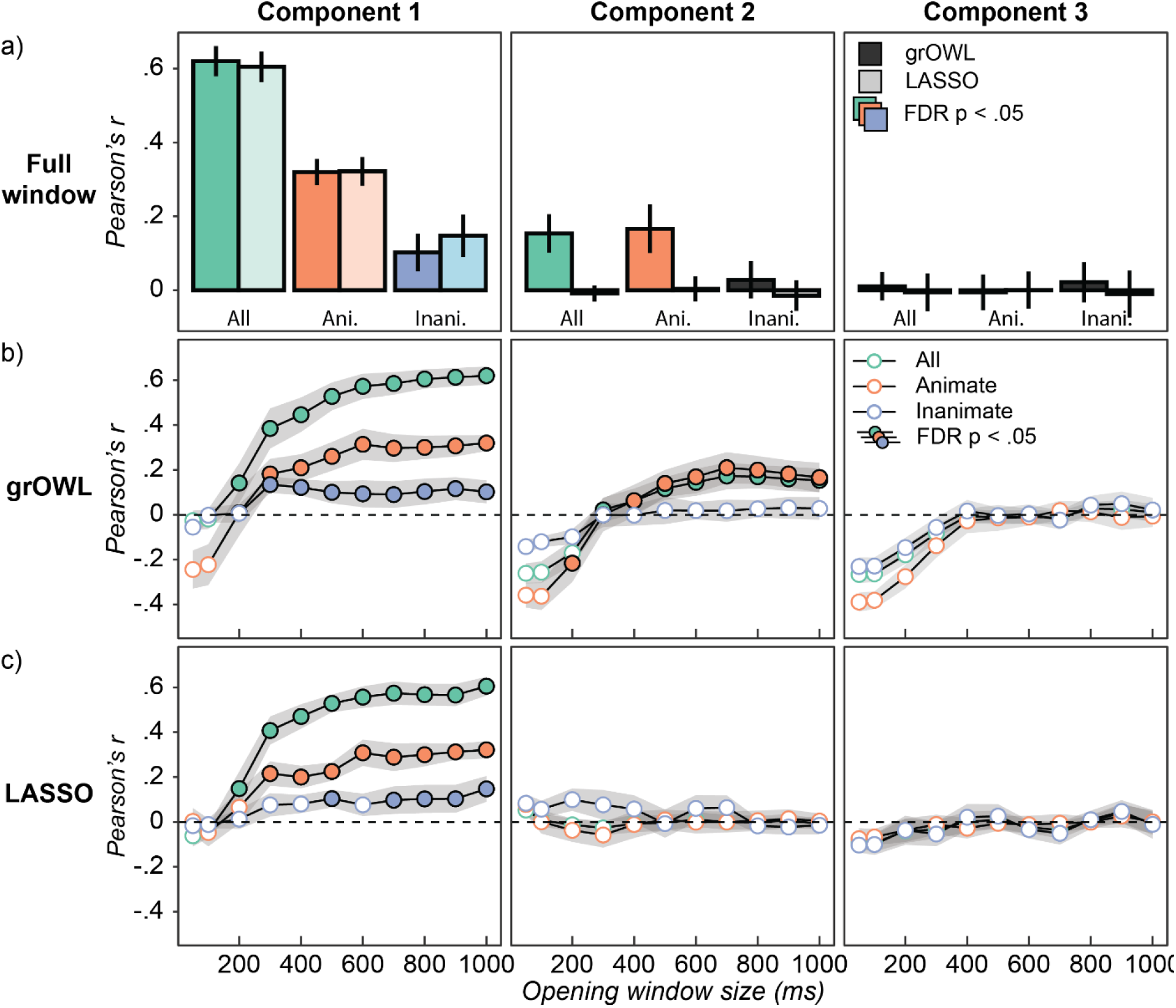
Uncentered ECoG decoding results. This figure is completely analogous to Figure 4 in the main paper, except that the raw correlations are presented, without centering on the mean of the permutation distribution. We see that the predictions of models fit with grOWL to small windows, including the least activation since stimulus onset, tend to be negatively correlated with the target values. This is due to the models being assigned very small weights, such that the variance in the predicted values associated with the training set means is large relative to the variance associated with the variability in the neural data. It is possible for negative correlations to be reliably higher than the expected value until the permutation-simulated null, as seen for the animate-subordinate structure along dimension 2 in the 200-ms window.

## Appendix D: Reconstructing item similarity from SVD coordinates

The main analysis uses the first three root-weighted singular vectors as the target for fitting decoding models with RSL. Predicted coordinates are then used to generate predicted semantic distances between pairs of held-out stimuli. To understand the possible causes of domain differences in prediction accuracy, we considered how well pairwise distances can be reconstructed from the first three singular vectors and values computed from the true matrix with a subset of items excluded. The goal was to assess whether the first three components capture inter-items distances equally well for both domains or whether such distances are better approximated for one domain than another.

To this end we conducted the following analysis. We generated 10,000 random splits of the semantic norming data, each consisting of 90 training items and ten test items (five animate and five inanimate) as in the ECoG study. For each, we computed the SVD on the semantic distance matrix for only the items in the training set. We then projected test-set items into the resulting space by matrix multiplying each item’s similarity with the training set items:

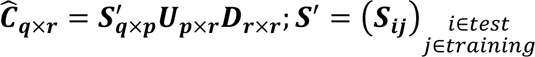

Here *q* = 10 is the number of test set items, *p* = 90 is the number of training-set items, *r* = 3 is the rank of the embedding, 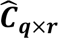 is a matrix containing estimates coordinates for the test items, ***S***′ is a matrix containing the true distances from test items to training items, and ***U*** and ***D*** are matrices containing the singular vectors and values, respectively, computed the training items. From the estimates coordinates in 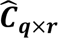 , the predicted similarity matrix can be obtained as 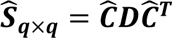. To measure how well the true distances are captures by the SVD, we computed correlations of lower-triangular values between ***S*** and 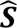, either over all values at once or correlating the animate-animate, inanimate-inanimate, and animate-inanimate similarity values separately.

**Figure S3.**
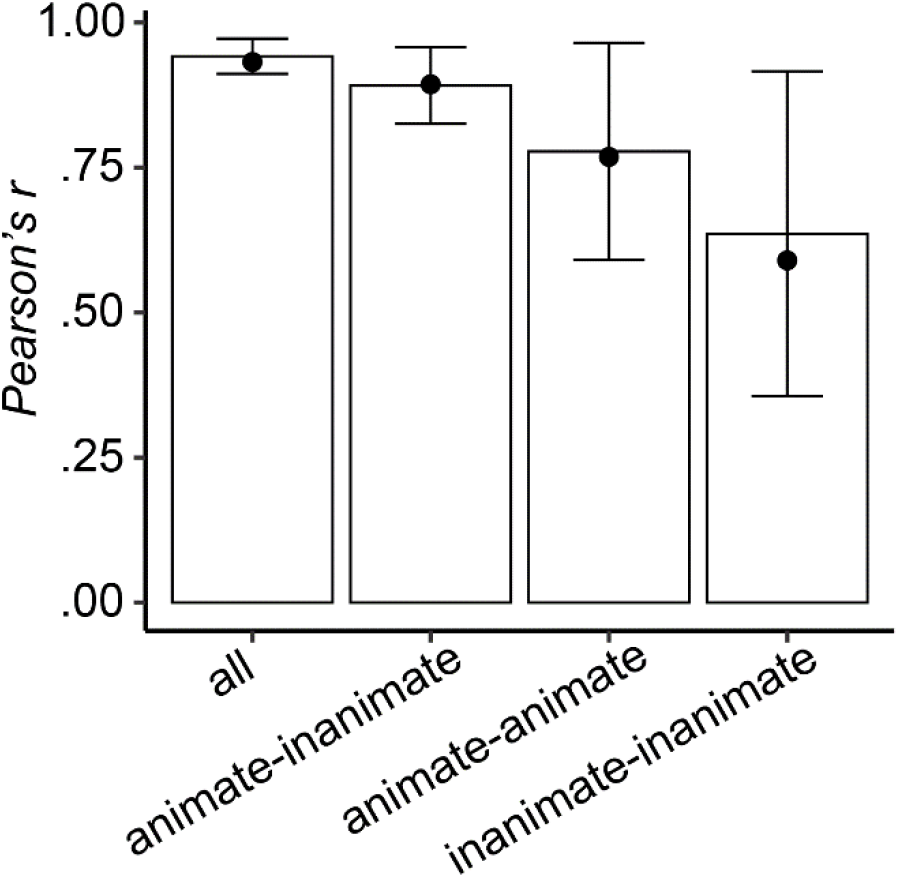
Correlation between true and reconstructed similarity matrices. Reconstructed similarity matrices are based on a SVD of a subset of the full similarity matrix expressing the pairwise similarities for 90 items, which is then used to reconstruct the similarities for the remaining 10 items as described in the text. This process was repeated 10,000 times with different 90/10 splits and each reconstruction was correlated with the corresponding true pairwise similarities. Bars show the average correlations over all similarities, just the between-category similarities (animate-inanimate), and just the within-category similarities (animate-animate and inanimate-inanimate). Error bars report the standard deviation of the means. The results for the specific 90/10 split used in the analysis of ECoG data reported in the main document are shown as black dots.

Figure S3 shows the average correlations for each selection of similarities over the 10,000 splits as bars with error bars reflecting the standard deviation over splits. The mean over the 10 cross-validation splits used in the ECoG study are overlaid as points. Predicted similarities that include or isolate between-category structure are most correlated with the true similarity structure. Predicted within-category similarities are less correlated, with predictions for inanimate items being less accurate than for animate items. In other words, a low-rank approximation of the semantic similarity matrix best captures between-domain distances and provides a better approximation of within-animate than within-inanimate semantic structure. Thus, decoding models face an intrinsic disadvantage at capturing within-domain structure generally, and within-inanimate structure specifically.

## Appendix E: Correlation between predicted and full-rank semantic distances

In evaluating how well the RSL decoding models predict pairwise semantic similarities, the main paper reports the correlation between predicted similarities and similarities reconstructed directly from the three singular vectors computed from the “true” full-rank matrix. We will refer to this as the *low-rank* similarity matrix. The low-rank similarity matrix represents the best possible prediction, given that only three components of the true matrix were used to fit the decoding model. Here we report correlations with the *full-rank* similarity matrix—the actual pairwise similarities amongst feature vectors as described in the main text. The low-rank and full-rank similarity matrices are very similar, *r* = .94, such that the low-rank similarity matrix accounts for ∼90% of the variance in the full-rank matrix. As one would expect given the high correlation, **Figure S4** and **Figure 5** from the main document show the same pattern of results. Filled circles indicate significance at Westfall-Young FWER corrected α = .05.

**Figure S4.**
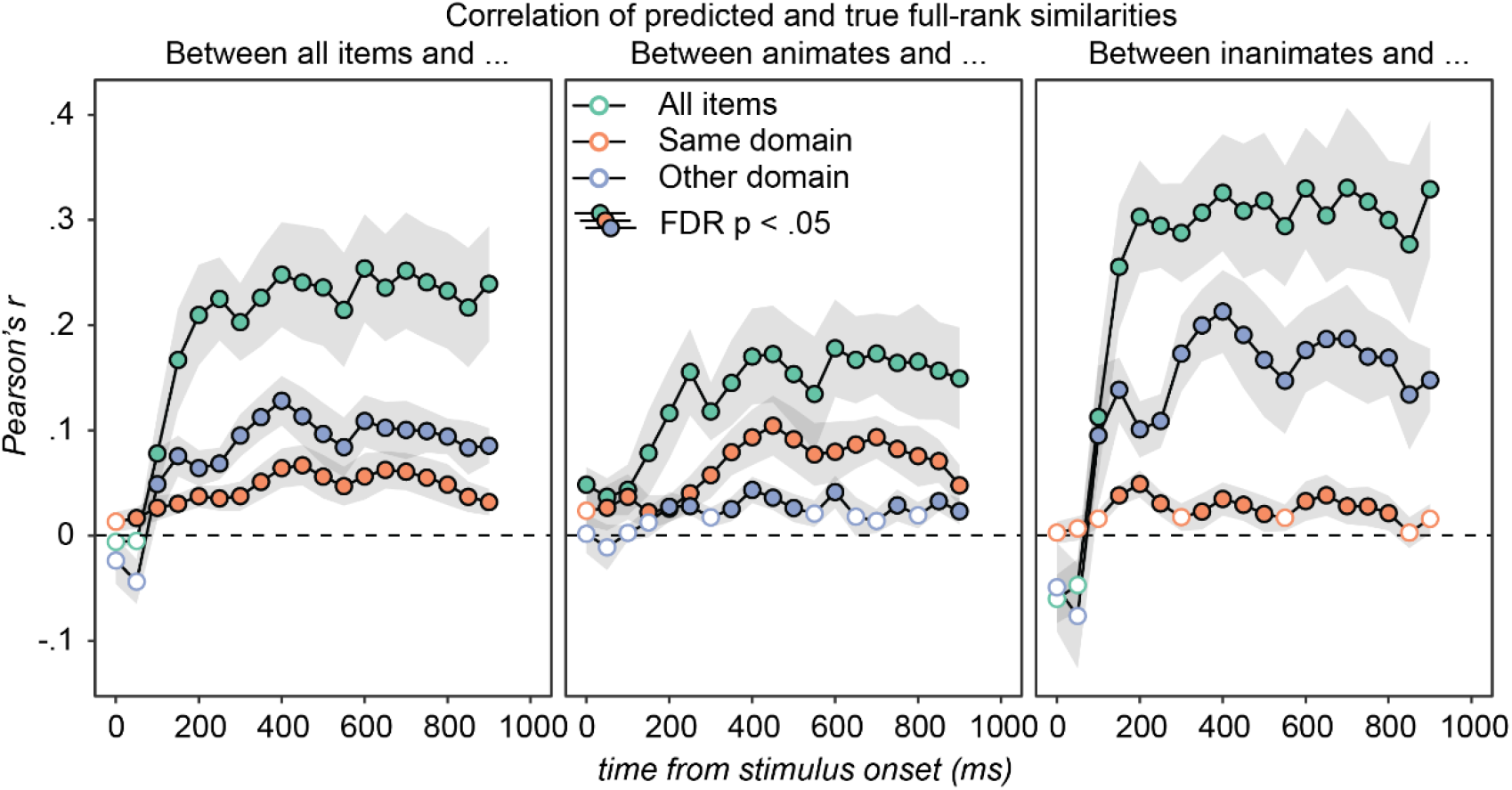
Correlation between predicted semantic similarities and *full-rank* cosine similarities as computed from the feature vectors. These results are very similar to **Figure 5** in the main document which involves the *low-rank* similarity matrix, as reconstructed from the three semantic dimensions modeled in the primary analysis. The results are so similar because the first three components account for ∼90% of the variance in the full-rank cosine similarity matrix. The interpretation is the same as **Figure 5.**

## Appendix F: Temporal generalization of models fit within moving windows

The model fit within a moving window can be used to decode target signal from the data within all other moving windows, forming a temporal generalization profile forwards and backwards through time. In Figure S5, each row in each panel is the temporal generalization profile for one 100ms window. Colored points are significant with FDR-corrected p<.05, correcting over the three panels in a single row (i.e., the multiple subsets relative to a shared dependent variable). Profiles are shown for RSA-style paired-matrix correlations (Cosine Similarities) and for correlations between model predictions and the target embedding along each semantic component. Each analysis can be performed over all items or by restricting to within-animate or within-inanimate similarities.

Generalization is better at later time points, but some degree of reliable generalization is seen through time for most models that perform well on the window it was fit within for component 1 and the aggregate cosine similarities. However, this is not seen when decoding components 2 and 3. Models fit early in the epoch do not generalize to later time points, and models fit late in the epoch do not generalize to early in the epoch.

The terms “early” and “late” are intentionally vague. We are not arguing for a qualitative, discrete shift in representational content at a particular point in time. If concepts encoded in the ATL follow a non-linear activation trajectory, as is often observed in deep network architectures, that is sufficient to make “early” and “late” representations importantly different while still conveying aspects of the target structure (Rogers et al., 2021).

**Figure S5.**
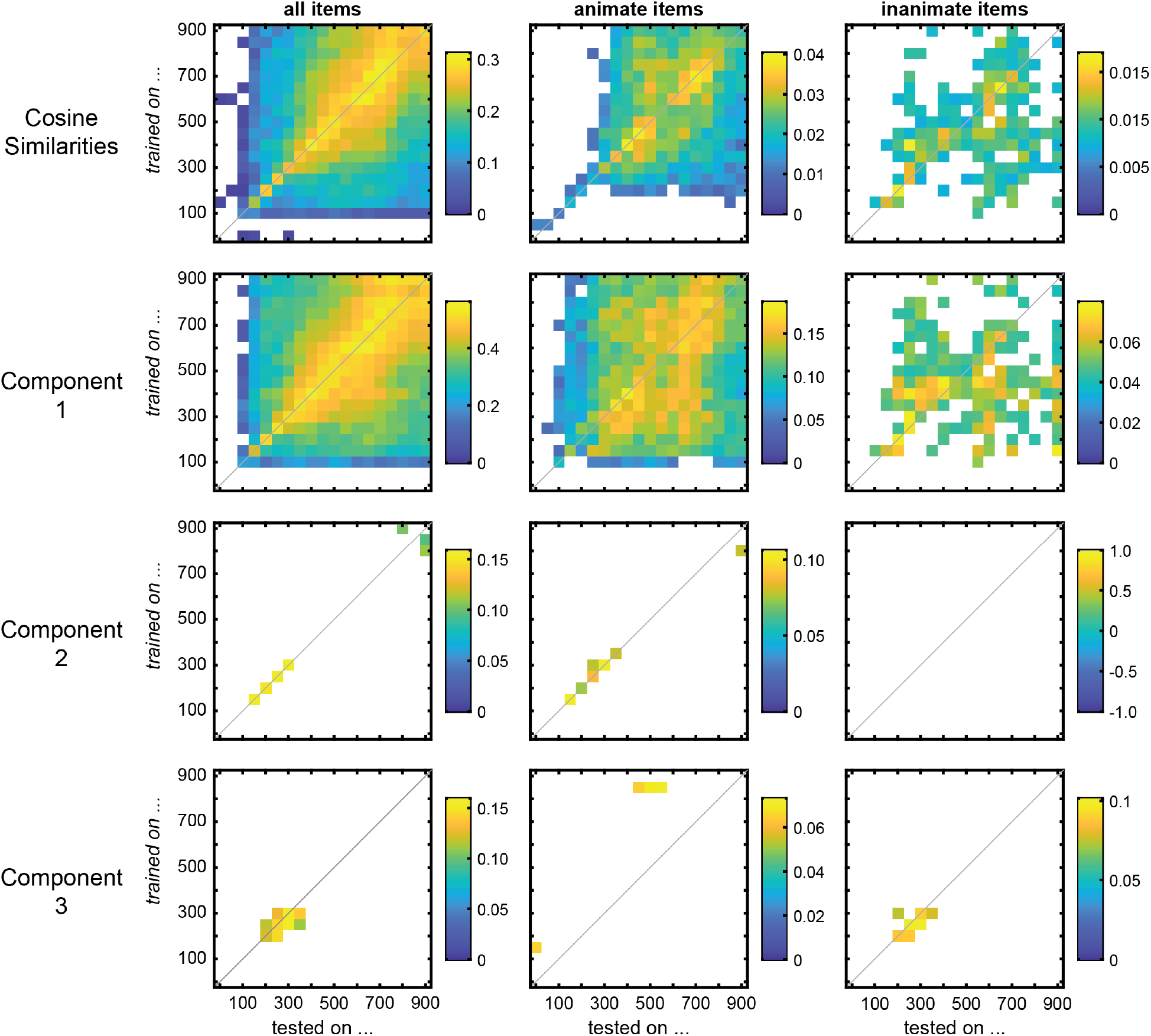
Temporal generalization profiles for each model fit within a single 100ms-moving-window and tested on all windows. Colored points are significant with FDR-corrected p<.05, correcting over the three panels in a single row (i.e., the multiple subsets relative to a shared dependent variable). The first row shows the correlations between predicted similarity matrices and the target similarity matrix. Subsequent rows show the correlation between single predicted and target components.

## Appendix G: Anatomical distribution of decoding model weights across vATL

The decoding results reported in the main paper demonstrate that multidimensional semantic structure can be decoded from LFPs recorded from the vATL, and that decoding the second semantic dimension depends on regularization via the grOWL. This implies that local neural populations do not independently encode each semantic dimension, but simultaneously express multiple orthogonal components of the target semantic matrix. To evaluate whether this is so, we examined the spatial distribution of weights across electrodes in the decoding models predicting variation on the first via the second semantic dimensions for opening windows of size 200, 400, 600, 800, and 1000ms in the eight patients with electrodes implanted in the left hemisphere. To obtain a single number at each electrode for each model (each component is associated with a distinct model), the weight magnitudes (absolute values) were summed over time. Because the number of electrodes and their placement varied across participants, we projected these sums onto a standard MNI cortical surface and smoothed along the surface with a 4mm Gaussian kernel before averaging across participants. Points on the surface that are associated with data from fewer than six participants are excluded from analysis (i.e., cortical areas outside the red-line contour shown on surface plots in Figure 6a). To determine which sums are larger than expected by chance, the data for each participant within each window was fit to 1,000 random row-wise permutations of the target structure. Each of these solutions were projected onto the MNI surface as described above, resulting in 1,000 surface maps per participant. Then we sampled a map from each participant to simulate one “experiment”, and the data were summed and averaged as above. This was repeated 10,000 times to obtain a group-level null distribution (Stelzer et al., 2013).

**Figure S6.**
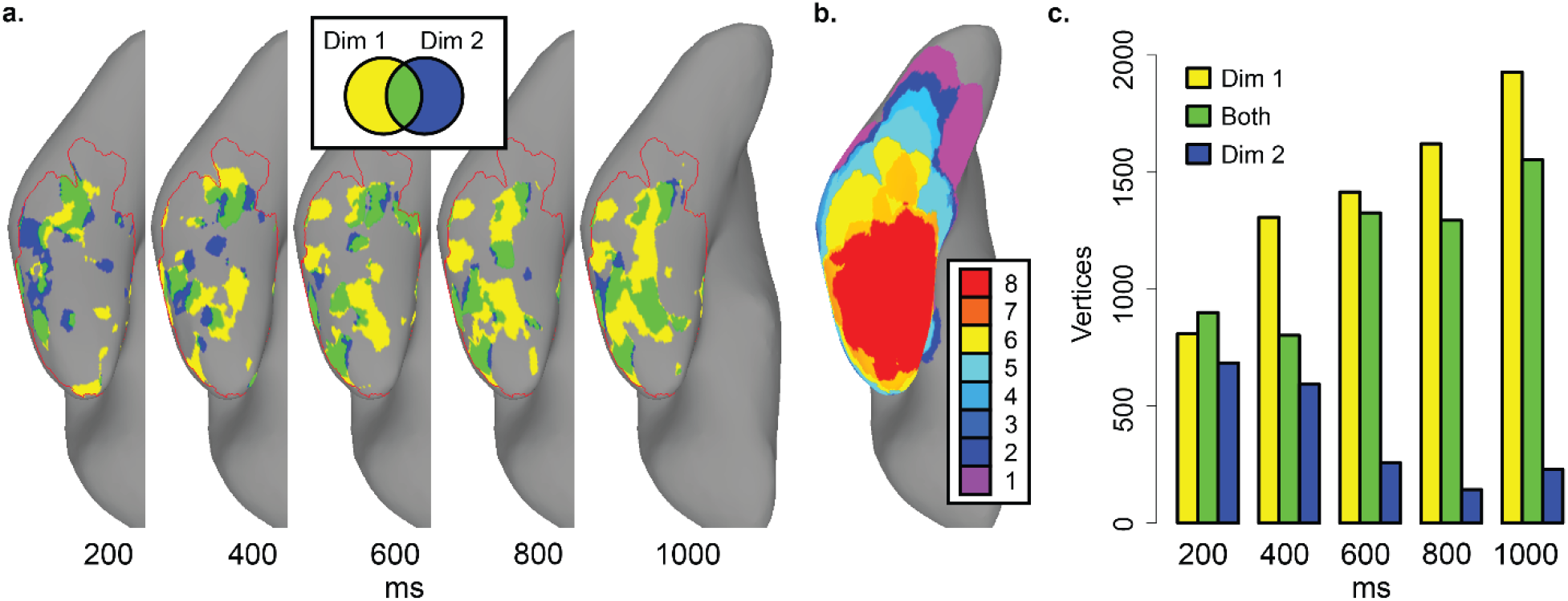
Significant vertices on the standard MNI N27 surface reconstruction displayed with AFNI SUMA for eight patients with left lateralized language (confirmed with the WADA test). a) RSL model coefficients (multi-task linear regression with grOWL regularization) associated with points in MNI space were projected onto and smoothed along the surface with a 4mm Gaussian kernel, separately for dimensions 1 and 2. This was repeated for 10,000 group-level maps simulating the null distribution at each vertex by permutation. Only points on the surface where data from 6 or more patients is available are considered; vertices meeting this criterion are contained within the red contour. Vertices with average weight magnitude larger than 9,500 of the 10,000 group-level values in the permutation distribution for that vertex are shown: points that meet these criteria for dimension 1 are shown in yellow, for dimension 2 are shown in blue, and for both are shown in green. b) Number of electrodes contributing to the data at each vertex after applying a 4mm Gaussian kernel. c) Tabulation of significant vertices.

Figure S6a displays the points on the cortical surface where the real average is larger than 9,500 of the values in this null distribution. Points on the surface that are associated with large model weights for the first component (but not the second) are shaded yellow; points that are associated with the second component (but not the first) are shown in blue. Points on the surface that are associated with large weights for both components in the same model are shown with green. Substantial overlap is observed in all windows, with overlap increasing with window length (**Figure S6c**). Both semantic dimensions are encoded across the vATL with little obvious functional localization, and the apparent contributions of a given area to each dimension appear to change depending on the size of the decoding window. The results are consistent with the view that many vATL regions contribute to both dimensions of semantic structure, though they do not rule out the possibility that there exist small local regions independently encoding each dimension scattered across cortex.

## Appendix H: Predicted embeddings

Representational similarity learning (RSL) involves decomposing a target representational similarity matrix (RSM) into orthogonal components (e.g., with singular value decomposition; SVD), composing a target embedding from a small selection of those components that explain most of the variance in the full matrix, and then training a linear model to decode each component in the target embedding. In our work the target embedding was three-dimensional (first three singular vectors of the RSM), and training and testing was performed with 10-fold cross validation such that every stimulus was a test item once. A model of the embedding determined with respect to the training set can be used to map spatiotemporal patterns of neural activity to predicted positions along each dimension of the embedding space (i.e., a coordinate referring to a point within the embedding space), including patterns associated with items assigned to the test set and therefore excluding from the training set. Test set predictions are obtained by weighting and then summing the neural features in the way that minimized the difference between predicted and target coordinates for *training set* items, and so reflect where the model expects these items to be located based on their neural pattern similarity with items in the training set.

By collecting these predicted coordinates for test set items across the 10 cross validation folds, we can visualize how each of the 100 items (50 animate, 50 inanimate) is positioned along each of the three target dimensions by models trained over different windows of time, averaged over the 10 participants. This is visualized in Figure S7. As expected, given the results reported in Figures 4 – 6 in the main document and the proportion of variance in the target RSM attributed to each dimension, separation of animate from inanimate items along the first dimension is most salient in these predicted embeddings. This separation progresses over the first 300ms and remains relatively stable thereafter. Categories do not clearly separate along dimensions 2 or 3; there is variance along this dimension that systematically correlates with the target structure (Figure 4), but this presentation of predicted embeddings is not designed to convey how within-category variance (e.g., among land animals) is correlated with each target dimension. That there is stretching along dimensions 2 and 3, but not clear category separation, may reflect the smaller effect size of the within-domain correlations we report in the main paper, but is also consistent with the within-domain variance decoded from the ECoG signal being graded and continuous within sub-categories, not driven by ever-more minute category fractionization.

**Figure S7.**
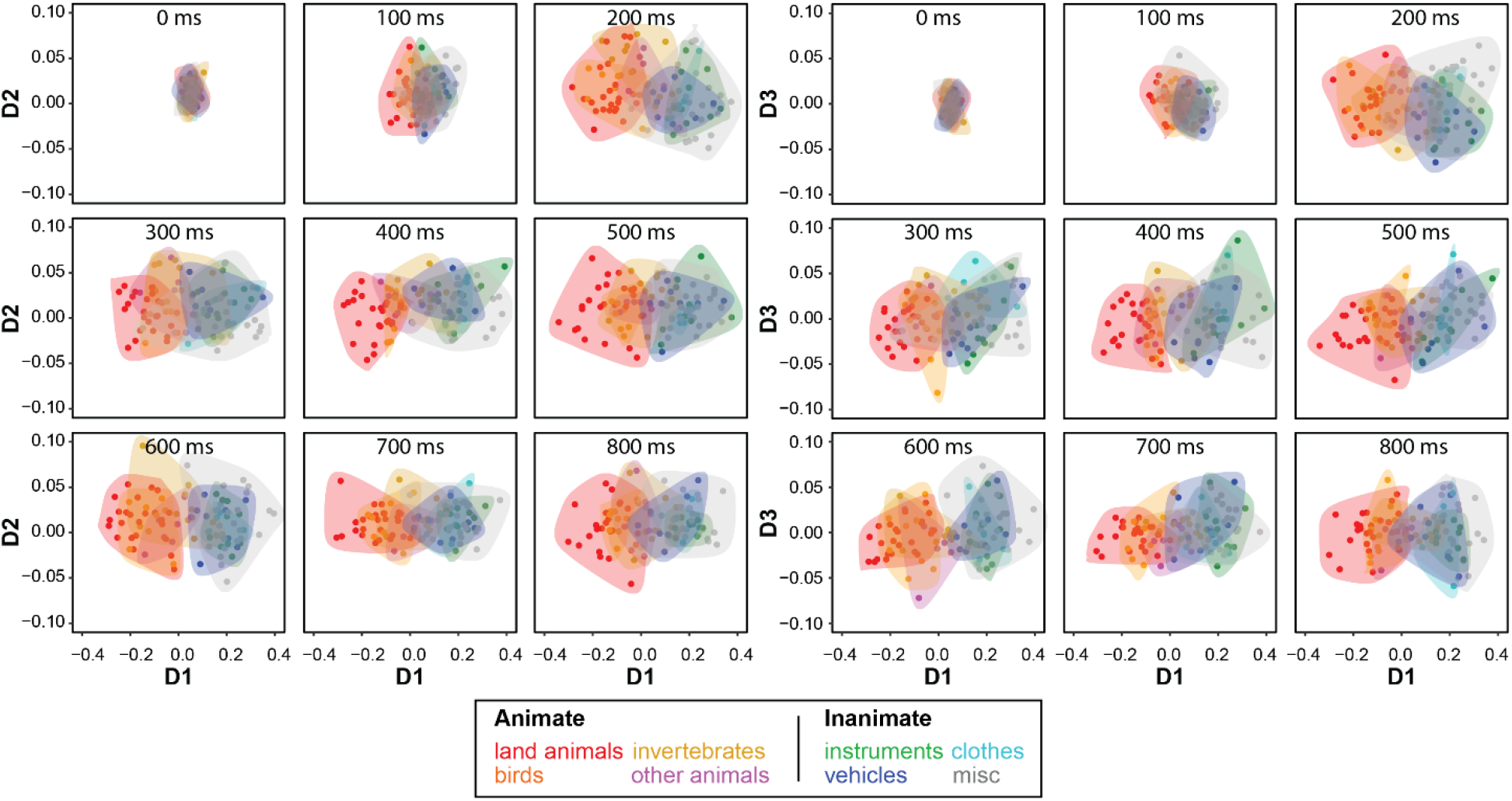
Predicted semantic embeddings for the 100 stimuli, averaged over 10 participants. Dots and shaded areas enclosing dots belonging to the same category are colored as in Figure 1. Which items belong to each category is reported in Appendix A, Table S1. Shaded areas were drawn by identifying the convex hull for the set of points and tracing the hull with a spline.

